# Midkine Attenuates Aβ Fibril Assembly and Amyloid Plaque Formation

**DOI:** 10.1101/2025.03.20.644383

**Authors:** Masihuz Zaman, Shu Yang, Ya Huang, Jay M. Yarbro, Yanhong Hao, Zhen Wang, Danting Liu, Kiara E. Harper, Hadeer Soliman, Alex Hemphill, Sarah Harvey, Shondra M. Pruett-Miller, Valerie Stewart, Ajay Singh Tanwar, Ravi Kalathur, Christy R. Grace, Martin Turk, Sagar Chittori, Yun Jiao, Zhiping Wu, Anthony A. High, Xusheng Wang, Geidy E. Serrano, Thomas G. Beach, Gang Yu, Yang Yang, Ping-Chung Chen, Junmin Peng

## Abstract

Proteomic profiling of Alzheimer’s disease (AD) brains has identified numerous understudied proteins, including midkine (MDK), that are highly upregulated and correlated with Aβ since the early disease stage, but their roles in disease progression are not fully understood. Here we present that MDK attenuates Aβ assembly and influences amyloid formation in the 5xFAD amyloidosis mouse model. MDK protein mitigates fibril formation of both Aβ40 and Aβ42 peptides in Thioflavin T fluorescence assay, circular dichroism, negative stain electron microscopy, and NMR analysis. Knockout of *Mdk* gene in 5xFAD increases amyloid formation and microglial activation. Further comprehensive mass spectrometry-based profiling of whole proteome and detergent-insoluble proteome in these mouse models indicates significant accumulation of Aβ and Aβ-correlated proteins, along with microglial components. Thus, our structural and mouse model studies reveal a protective role of MDK in counteracting amyloid pathology in Alzheimer’s disease.

---

Alzheimer’s disease (AD), an irreversible and progressive neurodegenerative disorder, is the primary cause of dementia^1^. Genetic studies have revealed disease-specific genes (APP, PS1, PS2), high-risk genes (APOE4, TREM2, UNC5C), and a large number of risk loci associated with amyloid, tau, endocytosis, and immunity^2–4^. The dysregulation in the AD proteome within the brain is characterized by the accumulation of Aβ amyloid plaques^5^, tau neurofibrillary tangles^6^, and other aggregated proteins, including TDP-43^7,8^ and U1 snRNP^9–13^. The recent FDA approval of lecanemab and donanemab^14,15^, both being amyloid-targeting antibodies, lend support to the amyloid cascade hypothesis. However, the limited clinical benefits for AD patients suggest the presence of alternative disease mechanisms and imply that intervention at a late disease stage may be insufficient.

Complementary to the genetic search for AD disease genes, we and others have performed large-scale proteomic profiling, uncovering consistent protein alterations in post-mortem AD brains across multiple large cohorts^16–23^. This has highlighted the deregulation of numerous molecular pathways and protein co-expression modules. Remarkably, a subgroup of proteins is highly upregulated and show a strong correlation with Aβ levels since the early prodromal stage throughout disease progression, including midkine (MDK), pleiotrophin (PTN), netrin 1 (NTN1), netrin 3 (NTN3) and the HTRA1 serine protease^16,19^. Interestingly, these protein changes occur independently of their mRNA transcript levels, indicating post-transcriptional regulation^16,19^, such as protein turnover ^24,25^. Moreover, these Aβ-correlated proteins have been found in the detergent-insoluble, aggregated proteome, and within amyloid plaques in AD brains, as confirmed by mass spectrometry-based profiling and immunohistochemical analyses^13,26–29^. However, the direct involvement of these Aβ-correlated proteins in plaque formation and AD pathogenesis remains to be elucidated.

The MDK/PTN family of cytokines, comprising only two members^30^, are among the most closely Aβ-correlated proteins identified in proteomic analyses^16,19^. They are multifunctional heparin-binding factors that promote growth, survival, and migration of various cell types, including neurons and immune cells^30^. In 1993, MDK was reported to localize within the amyloid plaques associated with AD^31^. Preliminary studies suggest that MDK might directly interact with Aβ, displaying high affinity and potentially mitigating Aβ’s toxic effects^32^, but the study had not received attention in the AD research field. Recent proteomic findings support MDK as a top candidate for modulating AD pathogenesis^16,19^, raising interest in fully exploring the role of MDK in AD pathogenesis.

In this study, we have thoroughly characterized the impact of MDK on Aβ assembly in various assays including Thioflavin T, circular dichroism, negative stain electron microscopy (EM), and NMR, demonstrating that MDK protein inhibits the formation of Aβ fibrils. Importantly, the *Mdk* gene knockout (KO) in the background of 5xFAD mice leads to the amyloid increase and microglial activation. Further proteomic profiling of these mouse models reveals the accumulated Aβ and related proteins in the brain of the 5xFAD/KO mice, supporting MDK’s protective role against amyloid pathology in AD.

## Results

### MDK, a top Aβ-correlated protein in human brain, attenuates Aβ fibril assembly

Although MDK was identified within the senile plaques of AD patients by antibodies last century^31^, its role in AD has not been well investigated. Recent unbiased mass spectrometry (MS)-based proteomic analyses have highlighted MDK as one of the top upregulated proteins highly correlated with Aβ^16,19^. To verify these findings, we re-analyzed an AD proteomic dataset comprising 111 human cases^16^, and performed Pearson correlation analyses of all identified proteins with Aβ and tau levels (**Supplementary Data 1**). Strikingly, among more than 8,400 identified proteins, MDK shows the best correlation with Aβ levels (R = 0.90) (**Fig. 1a**). In contrast, MDK displays a much weaker correlation with tau (R = 0.29), consistent with the observation that the levels of Aβ and tau do not strongly correlate in the human brain cases (R = 0.26). As a negative control, the abundant housekeeping protein ubiquitin (Ub) exhibits essentially no correlation with Aβ levels (R = 0.05). In the 5xFAD amyloidosis mouse model (referred to as FAD for simplicity), which overexpresses mutant human *APP* and *PSEN1* genes^33^, MDK accumulation was higher in FAD mice than in wild type (WT), as determined by western blotting of cortical brain lysates (**Fig. 1b**). Immunohistochemical (IHC) staining and immunofluorescent co-staining with Aβ confirmed the presence of MDK in plaque regions of both human brains (frontal cortex, **Fig. 1c, d**) and the FAD mice (cortex, **Fig. 1e, f**). These findings not only confirm the strong Aβ-MDK association in human but also suggest the FAD mice as a valuable model for studying the role of MDK in AD pathogenesis.

**Figure 1.**
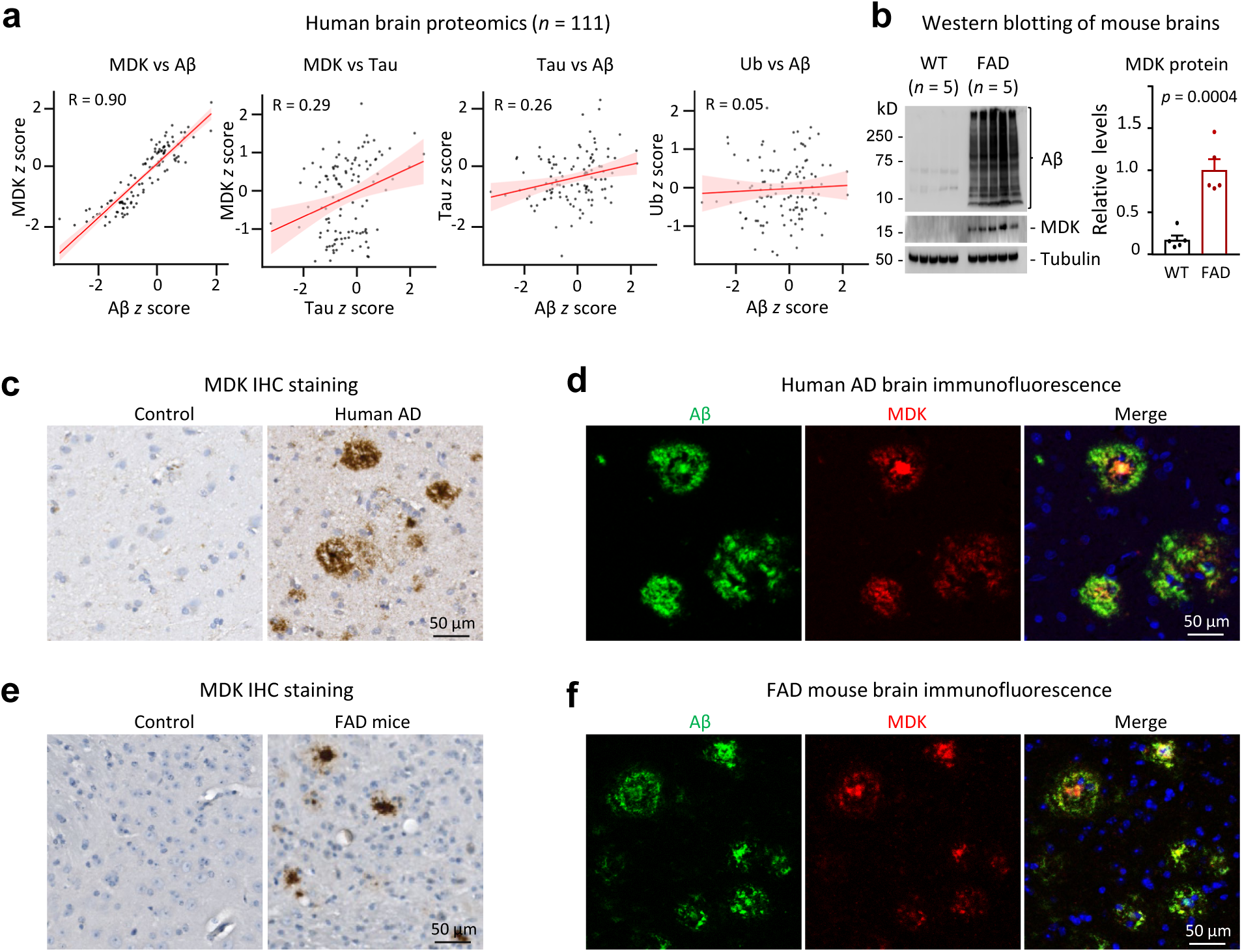
MDK is highly correlated and colocalized with Aβ in AD and in the FAD mouse model. **a**, Pearson correlation analysis on MDK-Aβ, MDK-tau, tau-Aβ and Ub-Aβ levels using human brain proteomic datasets (111 cases)^16^. Protein levels were standardized to z scores. **b,** MDK level is increased in FAD mouse brains, compared to WT (12-month-old, WT *n* = 5, FAD *n* = 5). Statistical significance is analyzed by two-tailed Student’s *t*-test. Data are shown as mean ± SEM. **c**, IHC staining of MDK in both control and AD cortical brain tissues, with the scale bar shown. **d**, Co-immunofluorescence staining of Aβ and MDK in AD cortical region, with the scale bar shown. **e**, IHC staining of MDK in WT and FAD cortical brain tissues, with the scale bar shown. **f,** Co-immunofluorescence staining of Aβ and MDK in FAD cortical region, with the scale bar shown.

Given previous reports that MDK protein directly binds to Aβ peptides^16,32^, we investigated its impact on Aβ fibrillization using recombinant human MDK in multiple biophysical assays. We purified and characterized MDK proteins expressed in mammalian 293 cells and *E. coli*. MDK is a secretory protein with its mature form containing five disulfide bonds (S-S, **Extended Data Fig. 1a, b**). The stained SDS gel indicated near homogeneity of recombinant MDK from either source after breaking S-S bonds (**Fig. 2a**). We then examined the formation of disulfide bonds in the proteins by both top-down and bottom-up mass spectrometry. As expected, the 293-expressed MDK proteins contain all five S-S bonds (**Fig. 2b, c**), whereas the bacteria-expressed version contains a mixture of three proteoforms with 5, 4, and 3 S-S bonds, with the 5 S-S bond form showing the highest abundance (**Fig. 2d**). The bottom-up analysis validated that all S-S bonds were formed with the anticipated Cys pairs (**Extended Data Fig. 1b, c**). We then evaluated the effect of recombinant MDK from both sources on the aggregation of synthetic Aβ40 and Aβ42 peptides. The fibrillization process was monitored by the fluorescence of thioflavin T (ThT), which increases upon binding to amyloid fibrils^34^. Although *E. coli*-expressed MDK contains mixed disulfide-bonded forms, it effectively inhibited Aβ fibrillization, similar to 293-expressed MDK, albeit with slightly less efficiency (**Fig. 2e, f, Extended Data Fig. 1d, e**). Given its comparable inhibitory effect, *E. coli*-expressed MDK was used for subsequent biophysical and structural studies due to its higher yield.

**Figure 2.**
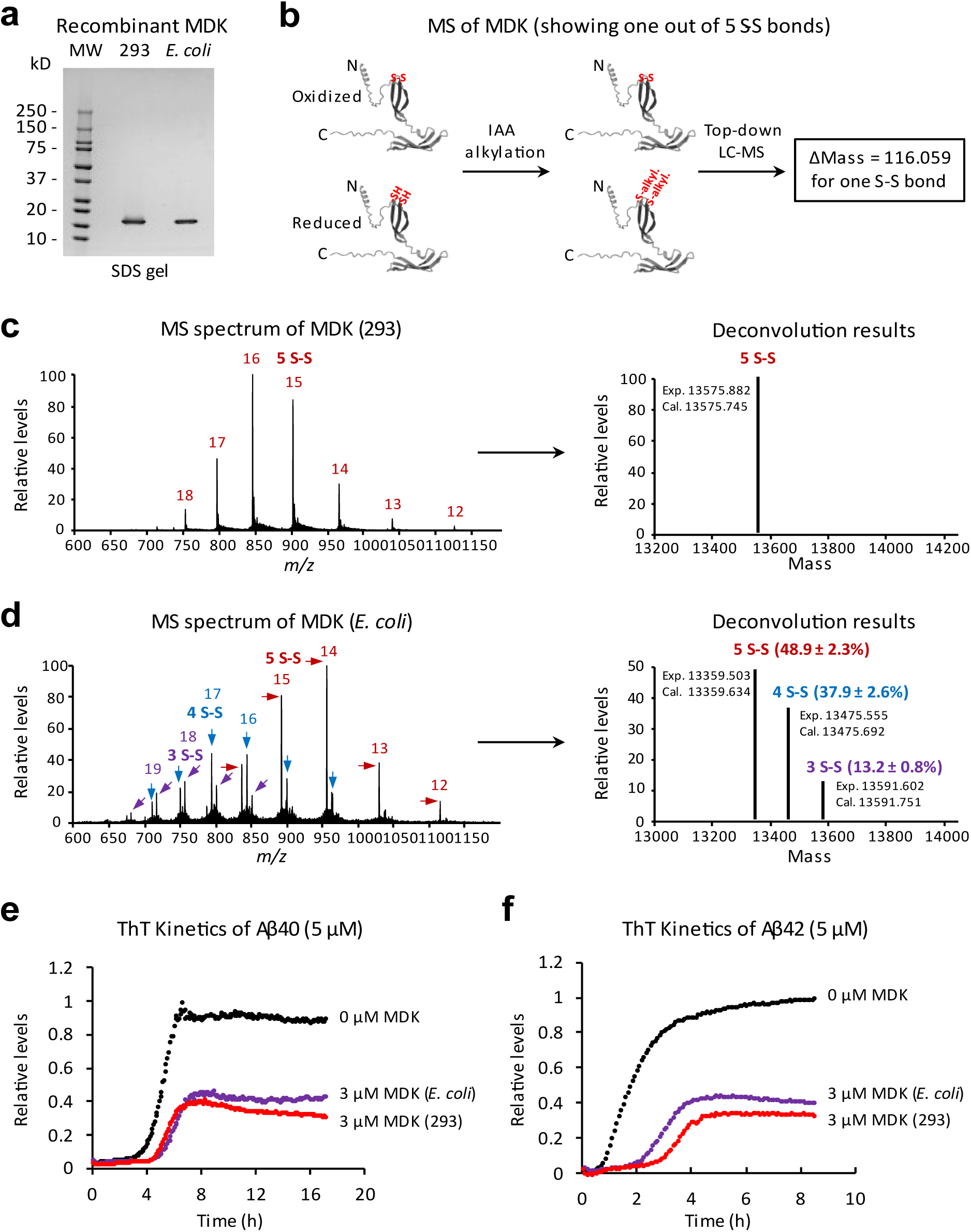
Characterization of recombinant MDK proteins and their impact on Aβ fibrillation. **a**, Highly purified recombinant MDK proteins expressed in mammalian 293 or *E. coli* cells, visualized on a stained SDS gel, with molecular weight (MW) markers. **b**, Schematic representation of top-down LC-MS for analyzing disulfide bond formation in MDK proteins. Free thiol (-SH) groups in Cys residues are alkylated by iodoacetamide (IAA), introducing a mass shift that differentiates them from proteins with intact disulfide bonds (S-S). The AlphaFold-predicted MDK structure (containing 5 S-S bonds) is downloaded, highlighting the N-terminus (N), C- terminus (C) and only one S-S bond (in red). **c**, Intact mass raw spectrum and deconvoluted results of 293-expressed MDK, revealing different charge states for a single protein form with 5 S-S bonds. **d**, Intact mass raw spectrum and deconvoluted results of *E. coli*-expressed MDK, showing charge states for three protein forms containing 5, 4, or 3 S-S bonds. The relative abundance of these forms was quantified from three independent scans, with data presented as mean ± SEM. **e-f**, Effect of 293- or *E. coli*-expressed MDK (3 µM) on Aβ40/42 (5 µM) fibrillation kinetics in ThT fluorescence assays. Results represent the average of 3 replicates.

We then fully characterized the aggregation propensity of Aβ peptides with recombinant MDK and the negative control protein Ub using thioflavin T assay, circular dichroism (CD) analysis, electron microscopy imaging, and Nuclear magnetic resonance (NMR) spectroscopy. When incubating Aβ40 peptide (5 µM) alone or with titrated MDK at concentrations ranging from 1 µM to 10 µM (**Fig. 3a**), Aβ40 displayed a time-dependent increase in fibrillization with a lag phase, growth phase and a final plateau. However, the fibrillization process was retarded by MDK with increased lag time and decreased ThT fluorescence intensity in a dose-dependent manner. We further used the AmyloFit software^35^ to determine the effects of MDK on microscopic steps during Aβ aggregation. The best fit revealed that MDK negatively impacts Aβ40 secondary nucleation rate constant (*k*_2_) and elongation constant (*k*_+_), as both log_2_(*k*_2_) and log_2_(*k*_+_) values showed a linear decrease with increasing MDK concentrations (**Fig. 3b**, **Extended Data Fig. 2a, b**). At low MDK concentrations (0.03 µM and 0.1 µM), ThT kinetics of Aβ40 remained largely unchanged, whereas at higher concentrations (1 µM and 3 µM), we observed consistent inhibitory effect (**Extended Data Fig. 2d**).

**Figure 3.**
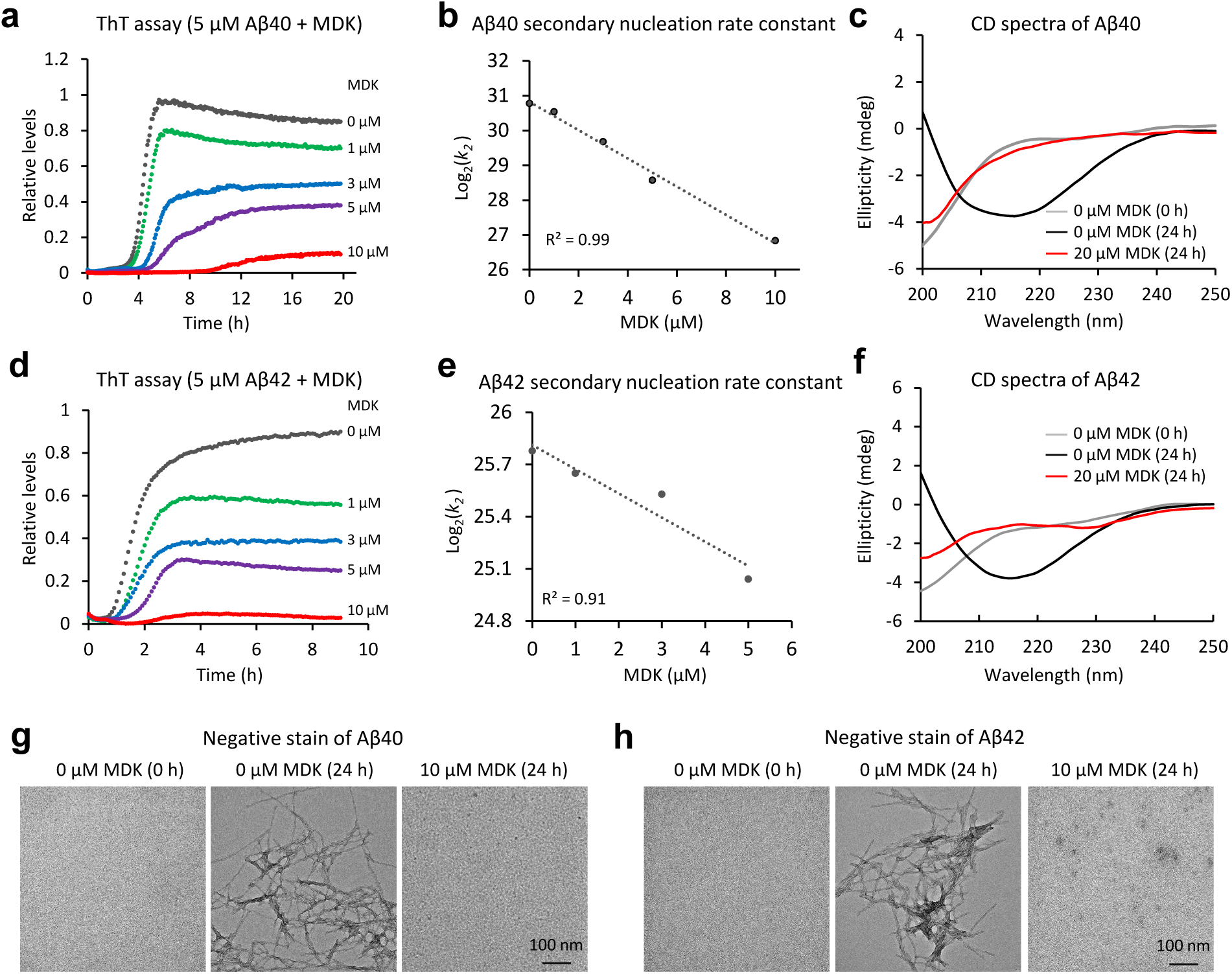
MDK inhibits the fibril assembly of Aβ40 and Aβ42 peptides. **a,** ThT fluorescence assay detecting Aβ40 amyloid fibril formation with titrated MDK protein (*n* = 3 replicates, averaged data shown). **b**, Analysis of secondary nucleation rate constant (*k*_2_) by globally fitting the ThT data using AmyloFit software^35^, estimating *k*_2_ at each MDK concentration in a unit (M ^-2^ h ^-1^), followed by log transformation and linear fitting of these values at different MDK concentrations. **c**, CD spectroscopy of Aβ40 (10 µM) with or without MDK, reporting ellipticity in millidegrees (mdeg). **d**, ThT assay of Aβ42 and MDK (*n* = 3 replicates, averaged data shown). **e**, *k*_2_ analysis for Aβ42 fibrillation using AmyloFit. Data points from the 10 µM MDK condition did not fit well and were excluded from the fitting. **f**, CD spectroscopy of Aβ42 (10 µM) with or without MDK. **g-h**, Negative stain EM of Aβ40/MDK and Aβ42/MDK samples, respectively, with the scale bars shown.

ThT fluorescence measurements are supported by far-UV CD spectroscopy, which detects the characteristic β-sheet structure of amyloid fibrils formed by peptide self-assembly. Consistently, CD analysis after 24 h incubation confirmed the inhibitory effect of MDK on fibrillization (**Fig. 3c**). Aβ40 alone transitioned from an unstructured random coil (0 h) to a β-sheet structure with a characteristic minimum around 218 nm (**Extended Data Fig. 2f**). In contrast, with the addition of MDK, Aβ40 largely retained its unstructured conformation, showing a negative peak near 200 nm, similar to Aβ40 monomers in a time course analysis (**Extended Data Fig. 2g**). We repeated all ThT and CD analyses using Aβ42 peptide, yielding similar results (**Fig. 3d, e, f, Extended Data Fig. 2c, e, h, i**). Electron microscopy imaging was used to detect the morphology of amyloid fibrils in Aβ-MDK samples. Amyloid fibrils were observed as end-point aggregates in samples of Aβ40/Aβ42 peptide (5 µM) alone (**Fig. 3g, h**). In contrast, fibril formation was abolished in the presence of high concentrations of MDK (10 µM, 2:1 molar ratio).

In negative control experiments, we validated highly purified Ub by an SDS gel and MS analysis (**Extended Data Fig. 3a, b, c, d**) and then performed ThT, CD and imaging experiments under similar conditions. Unlike MDK, Ub did not exhibit a noticeable inhibitory effect on Aβ aggregation (**Extended Data Fig. 3e, f, g, h, i, j**). Moreover, it is important to note that neither MDK nor Ub alone fibrillated under the same conditions, as confirmed by ThT binding assay, far-UV CD, and negative EM staining (**Extended Data Fig. 4a, b, c, d ,e, f)**. Together, these results indicate that midkine effectively inhibits the formation of amyloid fibrils *in vitro*.

In spite of the inhibitory effect of MDK on Aβ aggregation, we observed that Aβ could still assemble as fibrils in the presence of lower MDK concentrations, such as a 1:1 molar ratio, and subsequently examined the presence of Aβ42 fibrils under this condition using negative stain (**Extended Data Fig. 5a**). As recombinant Aβ fibrils assemble differently from endogenous Aβ filaments found in human brains^36–40^, we further investigated the presence of MDK in Aβ fibrils extracted from human AD brains. Immunogold negative stain EM with the MDK antibodies revealed that MDK molecules were clustered around the Aβ fibrils (**Extended Data Fig. 5b**). Moreover, we utilized the AlphaFold-multimer prediction tool^41^ to hypothesize possible Aβ-MDK interacting interfaces, suggesting that the N-terminus of Aβ (residues 1-12) might interact with the C-terminus of MDK (residues 74-80 and 84-89) (**Extended Data Fig. 5c, d**).

To further confirm the effect of MDK on Aβ fibrillization, we acquired ^1^H-^15^N HSQC NMR spectra of ^15^N-labeled monomeric Aβ40 (10 μM) with MDK titration (0, 5, and 10 μM, **Fig. 4a**). Reference spectra were acquired prior to incubation, with amide cross peaks assigned to specific residues^42^. After 48 h, a notable disappearance of cross peaks suggested Aβ40 peptide fibrillization. Remarkably, with MDK co-incubation at varying concentrations, the cross peaks re-appeared although at reduced intensities. As the HSQC cross peak intensities correlate with monomeric Aβ concentration^43^, we quantified the cross peak intensities for each residue (**Fig. 4b**), and calculated the averaged percentage relative to the pre-incubation reference of Aβ40. The MDK addition (5 μM and 10 μM) rescued the cross-peak intensities from 0.9±0.2% to 38.3±1.1% and 38.1±0.9%, respectively (**Fig. 4c**). We also observed a similar impact of MDK on Aβ42 fibrillization using a protocol of 24 h incubation (**Fig. 4d**). However, the same concentrations of MDK were less effective in recovering Aβ42 cross peaks compared to Aβ40 (**Fig. 4e, f**). For example, the MDK addition (5 μM and 10 μM) retrieved the cross-peak intensities from 4.8±0.5% to 12.8±0.8% and 21.8±0.8%, respectively. This aligns with our ThT, CD and negative stain EM data, supporting the notion that Aβ42 is more aggregation prone than Aβ40^39^. In contrast, ubiquitin (10 µM) used as a control did not restore the Aβ40/42 signal under similar experimental conditions, supporting the specificity of MDK’s inhibitory effect (**Extended Data 6a, b, c, d**). In summary, our comprehensive biophysical characterization strongly supports that MDK can modulate Aβ fibrillization *in vitro*.

**Figure 4.**
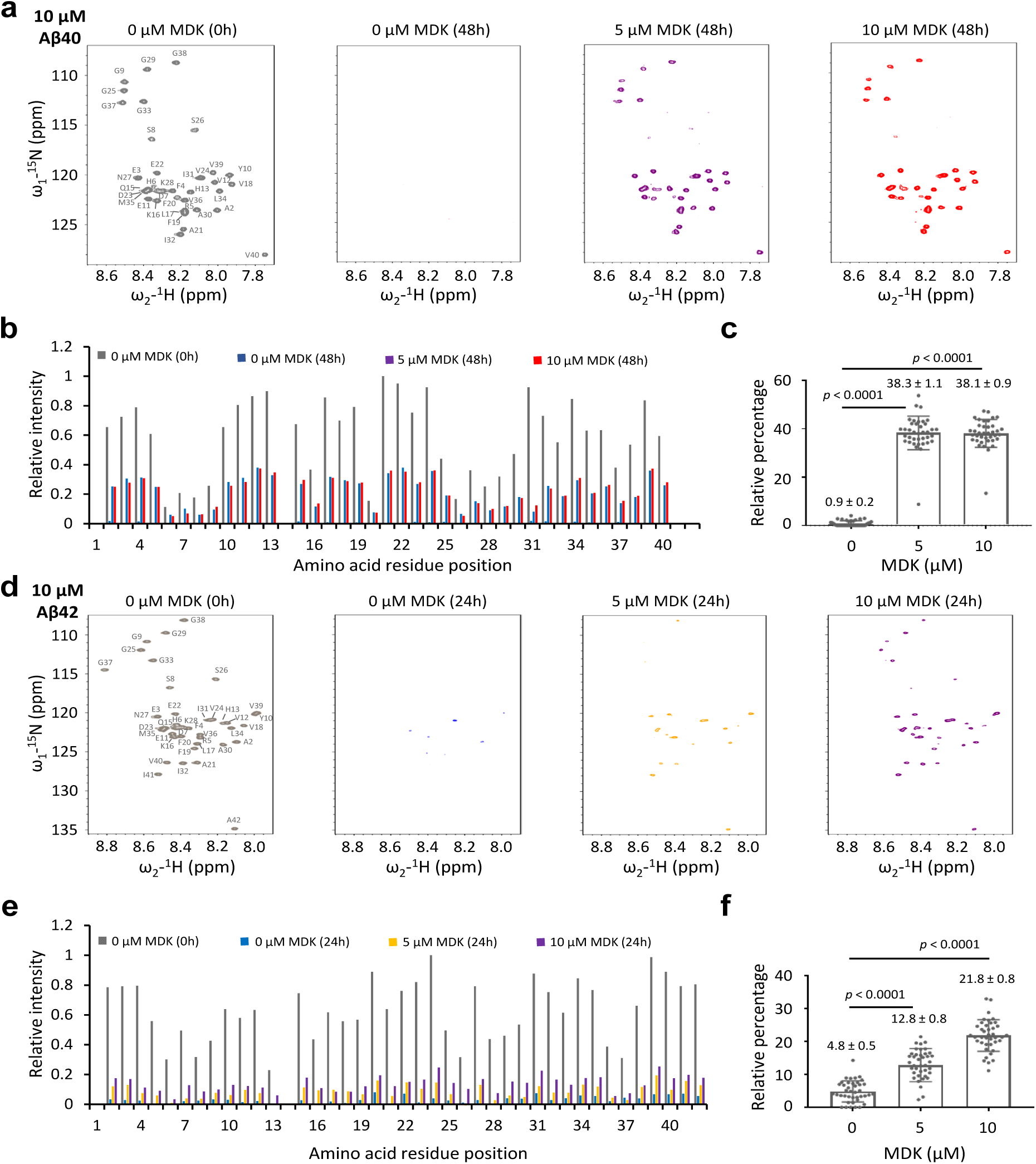
MDK rescues NMR signals of Aβ peptides. **a,** ^1^H-^15^N HSQC spectra of Aβ40 (10 μM) with MDK at 0, 5, or 10 μM in 50 mM Tris buffer (pH 7.5). Spectra were collected before incubation (left panel) and after 48 h incubation (right three panels). **b,** Relative cross-peak intensities for each residue, excluding D1 and H14. Intensities were normalized to a maximum value of 1. **c,** Relative percentage of Aβ40 intensities after 48 h incubation compared to pre-incubation values. Percentages were calculated pre residue and averaged, with data shown as mean ± SEM. Statistical significance was analyzed by two-tailed Student’s *t*-test. **d-f,** HSQC spectra of Aβ42 under similar conditions as Aβ40, but with 24 h incubation. Statistical significance was analyzed by two-tailed Student’s *t*-test.

### Genetic deletion of *Mdk* gene in FAD mice increases amyloid plaques and microglial activation

To investigate the role of MDK in Aβ fibrillization *in vivo*, we generated a mouse model with an *Mdk* gene knockout by CRISPR-mediated deletion of 23 base pairs in exon 3 of the mouse genome (**Fig. 5a**). The deletion was confirmed through genome sequencing and PCR analysis (**Fig. 5a**), with the absence of MDK protein verified by western blot analysis (**Fig. 5b**) and mass spectrometry (**Extended Data Fig. 7a, b**). In WT mouse brain, *Mdk* gene is mainly expressed from astrocytes and oligodendrocyte progenitor cells based on published single-cell RNA sequencing analysis^44^ (**Extended Data Fig. 7c**). We crossbred this knockout line with the FAD mice to produce offspring that were FAD mice with or without the *Mdk* knockout. The *Mdk* knockout resulted in a marked increase in Aβ levels in the cortices of 12-month-old mice brains, as shown by western blot analysis (**Fig. 5b**). ELISA analysis also indicated elevated accumulation of Aβ40 and Aβ42 in both Sarkosyl-soluble and -insoluble fractions of mouse cortices in the FAD/KO mice (**Fig. 5c, d**), whereas the ratio of Aβ40/Aβ42 was not significant changed (**Extended Data Fig. 7d**). These biochemical findings indicate that *Mdk* gene knockout contributes to the increased accumulation of two major Aβ species in the FAD genetic background.

**Figure 5.**
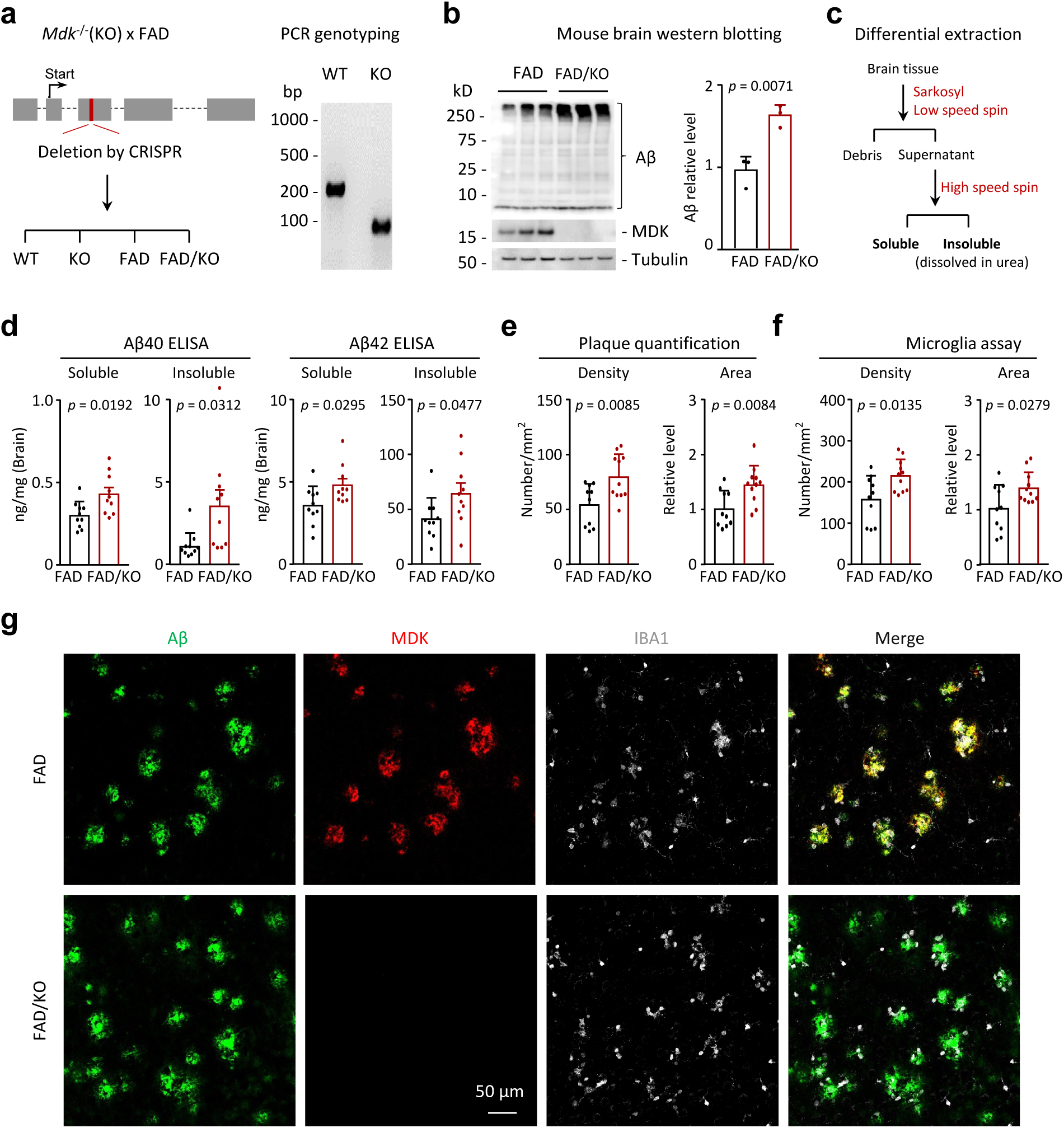
*Mdk* gene knockout in FAD mice results in Aβ accumulation, plaque increase and microglia activation. **a**, The diagram of the *Mdk* gene knockout and PCR genotyping of the WT and homogenous *Mdk* KO mice. The *Mdk* gene consists of five exons, including the first exon as the 5’ untranslated region. A CRISPR-mediated deletion was introduced within exon 3, disrupting the open reading frame and resulting in a gene knockout. Crossbreeding of *Mdk* KO with FAD mice generated four genotypes for comparison. **b**, Western blot analysis confirming the loss of MDK protein and increased Aβ in the brain of heterogenous FAD mice with homogenous *Mdk* KO (12-month-old, *n* = 3 replicates). The Aβ intensity was quantified from the western blot image and normalized by setting the mean value of FAD mice to 1. **c**, Differential extraction to generate Sarkosyl-soluble and -insoluble fractions of mouse brains. **d**, ELISA analysis of Aβ40 and Aβ42 in the Sarkosyl-soluble and -insoluble fractions of mouse cortices (*n* = 10 per group), with the unit of ng protein per mg brain tissue. The Sarkosyl-insoluble fractions were further dissolved in an 8 M urea-containing buffer and diluted for ELISA. **e**, Quantification of X34-positive amyloid plaque in the cortices (FAD *n* = 10, FAD/KO *n* = 11). The relative levels of area were calculated as the percentage of the total plaque area within the entire cortical region and normalized by setting the mean value of FAD mice to 1. **f**, Quantification of microglia density and area in the cortices (FAD *n* = 10, FAD/KO *n* = 11). **g**, Co-immunofluorescence staining of amyloid plaque, MDK and IBA1 in the mouse cortices. Scale bar, 50 µm. Statistical significance was analyzed using a two-tailed Student’s *t*-test if the two groups had equal variances, or by Welch’s *t*-test if the two groups had unequal variances. Data are shown as mean ± SEM. in c-f. Each point represents a data point of one mouse. Full statistical information is in Source Data Statistics.

Next, we characterized the FAD/KO mice through immunostaining to evaluate the effects of *Mdk* deletion on plaque formation and microglial activation in the brain. *Mdk* knockout led to increased plaque density and area, as detected by X34 staining, a fluorescent dye that binds to β-sheet structures in amyloid plaques^45^, in the cortices of 12-month-old mouse brains of both male and female mice (**Fig. 5e, Extended Data Fig. 7e, f, g**). This was accompanied by heightened microglial activation, evidenced by increased microglial density and area (**Fig. 5f**). In subsequent immunofluorescence staining studies, amyloid plaques detected by the Aβ antibody were found to highly colocalize with MDK protein in FAD mice, and the MDK staining was absent in the knockout mice (**Fig. 5g**). Additional co-staining with IBA1, a protein marker for microglia^46,47^, revealed a significant accumulation of microglia around the plaque areas (**Fig. 5g**), in agreement with the observed increase in microglial density and area in the FAD/KO mice. These results indicate that *Mdk* gene knockout leads to enhanced plaque formation and microglial activation in the FAD mouse model.

To understand the impact of *Mdk* knockout in the FAD mice at the molecular level, we analyzed the cortical brain proteome across four genotypes: WT (*n* = 8), KO (*n* = 8), FAD (*n* = 15), and FAD/KO (*n* = 17) as shown in **Fig. 6a**, Employing a refined tandem-mass-tag coupled with two-dimensional liquid chromatography and tandem mass spectrometry (TMT-LC/LC-MS/MS) platform^13,48–50^, we identified 154,013 peptides, corresponding to 9,441 protein groups (9441 genes) across all these samples with protein false discovery rate (FDR) below 0.01 (**Supplementary Data 2-3**). After loading bias correction (**Extended Data Fig. 8a**), principal component analysis (PCA) and clustering analysis clearly separate distinct mouse groups, indicating the high quality of the proteomic data (**Fig. 6b, Extended Data Fig. 8b**). As a positive control, MDK protein levels were elevated in FAD mice compared to WT and reduced to noise levels in knockout mice (**Fig. 6c**). In FAD mice, *Mdk* deletion led to a significant increase of Aβ peptide levels (**Fig. 6c**), but did not affect the expression level of the full-length APP protein (**Extended Data Fig. 8i, j**).

**Figure 6.**
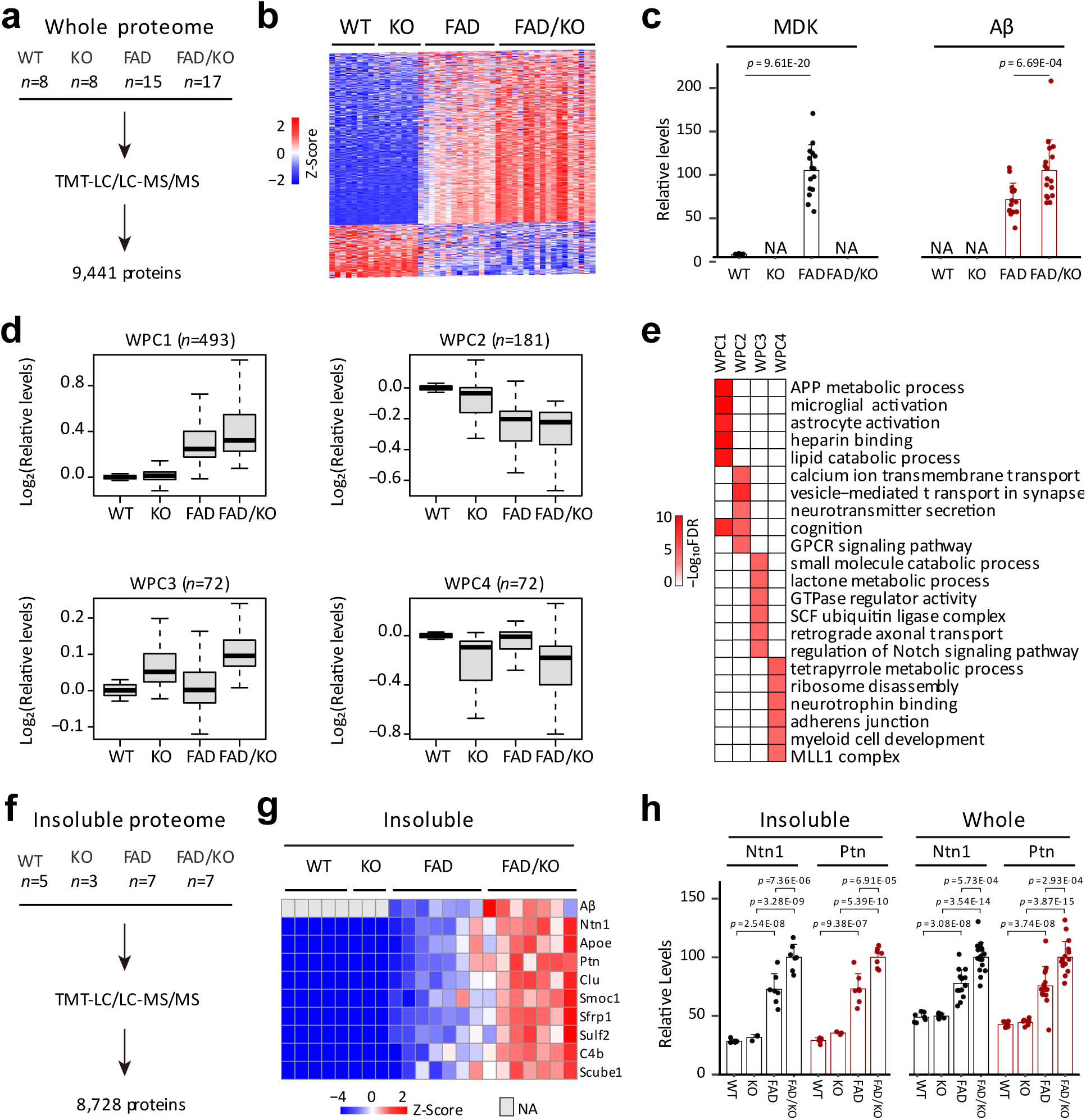
Brain tissue proteomics reveals that *Mdk* knockout leads to the accumulation of Aβ and Aβ-correlated proteins, along with microglia activation in FAD. **a**, Workflow for whole proteome analysis of the brain cortical region from four mouse genotypes (12-month-old, WT *n* = 8, KO *n* = 8, FAD *n* = 15, FAD/KO *n* = 17). **b**, Clustering of the whole proteome using DEPs to show genotype-specific grouping. **c**, MDK and Aβ levels extracted from the whole proteome analysis. **d**, Four major DEP clusters identified by WGCNA. Protein intensities were log transformed followed by Z-score normalization. **e**, Enriched pathways in the WPC proteins, identified by Fisher’s exact test. **f**, Workflow for insoluble proteome profiling (12-month-old, WT *n* = 5, KO *n* = 3, FAD *n* = 7, FAD/KO *n* = 7). **g**, Heatmap of selected proteins in the insoluble proteome, normalization by z score transformation. **h**, Ntn1 and Ptn protein levels extracted from the whole and insoluble proteome analyses. The two proteins were selected because they are highly correlated with Aβ levels in AD^16^, similar to MDK (**Supplementary Data 1**). Statistical significance was determined by a two-tailed Student’s *t*-test, with data presented as mean ± SEM.

To identify the differentially expressed proteins (DEPs) among the four genotypes, we performed an ANOVA analysis using a previously reported method^16^ with two criteria: (i) an FDR value of less than 5%, and (ii) a log2 fold change (FC) greater than twice the standard deviation (2 SD) in any of the three pairwise comparisons: WT vs KO, FAD vs WT, or FAD/KO vs FAD (**Extended Data Fig. 8c**). Following this analysis, a total of 818 DEPs were accepted (**Supplementary Data 4**). We then classified these DEPs into four whole proteome clusters (WPCs) using the weighted gene co-expression network analysis (WGCNA)^51^, followed by pathway enrichment analysis (**Fig. 6d, Supplementary Data 4-5**). The WGCNA program identified clusters of highly correlated proteins based on the DEP data by calculating pairwise correlations to construct a network, and applying a thresholding approach to emphasize strong correlation patterns. In detail, WPC1 (*n* = 493) exhibits a gradual increasing trend across the four genotypes from WT, KO, FAD to FAD/KO. This cluster includes Aβ and Aβ-correlated proteins^16^, such as Ntn1, Ptn, and Clu, as well as some marker proteins for microglia and astrocytes: Trem2, Ctss, Olfml3, Csf1r, Gfap, Aqp4, Pla2g7, and Gja1^52,53^ (**Extended Data Fig. 8d**), and it is enriched in four representative pathways: APP metabolic process, microglia activation, astrocyte activation, heparin binding and lipid catabolic process (**Fig. 6e, Supplementary Data 6**). In contrast, WPC2 (*n* = 181) shows notable decrease along the four genotypes, related to the pathways of calcium ion transmembrane transport, vesicle-mediated transport in synapse, neurotransmitter secretion, cognition, and G protein-coupled glutamate receptor signaling. Interestingly, this cluster includes the AD risk gene/protein Bin1 that regulates calcium homeostasis and neuronal excitability^54,55^.

In addition, both WPC3 (*n* = 72) and WPC4 (*n* = 72) display effects of *Mdk* knockout that are independent of the FAD genetic background. Proteins in WPC3 increase in *Mdk* KO mice and are enriched in six major pathways: small molecule catabolic process, lactone metabolic process, GTPase regulator activity, SCF ubiquitin ligase complex, retrograde axonal transport and regulation of Notch signaling pathway (**Fig. 6e, Supplementary Data 6**). For instance, Slc18a3 (the vesicular acetylcholine transporter) functions to transport acetylcholine into synaptic vesicles during cholinergic neurotransmission, regulating cognitive functions such as learning and memory, motor coordination, and synaptic plasticity^56^. WPC4 primarily consists of proteins that are significantly downregulated due to *Mdk* knockout, enriched in the pathways of tetrapyrrole metabolic process, ribosome disassembly, neurotrophin binding, adherens junction, myeloid cell development and MLL1 complex. For example, Nbr1 is a selective autophagy receptor that recognizes ubiquitinated proteins and damaged organelles, promoting their degradation through the autophagy-lysosome pathway^57^. Overall, the protein alterations found in WPC3 and WPC4 likely reflect the developmental function of midkine, implicating a pleiotropic impact of *Mdk* knockout. Nevertheless, within WPC1, we observed a distinct increase in Aβ and Aβ-correlated proteins, along with the activation of microglia and astrocytes, supporting that *Mdk* knockout exacerbates pathological severity.

Furthermore, we validated the enhanced pathology in FAD/KO mice by profiling the detergent-insoluble proteome, which is enriched in aggregated proteins, across the four mouse genotypes. A total of 169,864 peptides from 8,728 unique protein groups (8,728 genes, **Fig. 6f, Supplementary Data 7**) were identified. After performing quality control analyses, such as loading bias normalization and PCA (**Extended Data Fig. 8e, f**), we found that 314 DEPs were elevated in the insoluble proteome of FAD/KO mice compared to FAD mice (**Extended Data Fig. 8g, Supplementary Data 7**). Remarkably, the FAD/KO mice showed an accumulation of Aβ and other plaque-associated proteins in the insoluble proteome relative to FAD mice. A heatmap highlights some of these proteins, consistently detected in both the whole and insoluble proteomes, including Ntn1, Apoe, Ptn, Clu, Smoc1, Sfrp1, Sulf2, C4b and Scube1 (**Fig. 6g, h, Extended Data Fig. 8h**). It should be noted that the expression of full-length APP in these mouse lines was not affected by *Mdk* gene knockout in the whole proteome analysis (**Extended Data Fig. 8i, j**).

In summary, the knockout of *Mdk* gene within the FAD mice resulted in an increase in Aβ and Aβ- associated proteins, amyloid plaque formation, and microglial activation. This was substantiated through experiments of western blotting, ELISA, immunostaining of brain tissues, and extensive proteomic profiling of both the whole and detergent-insoluble proteomes in the brain.

## Discussion

Midkine is one of the most correlated proteins with Aβ peptides in the proteomic analysis of post-mortem human brain tissues of Alzheimer’s disease. Our biophysical and NMR studies show that MDK inhibits Aβ assembly and fibril formation *in vitro*. Consistently, functional *in vivo* analysis in the FAD mouse model shows that CRISPR-mediated *Mdk* knockout leads to increased amyloid deposition and concurrent microglial activation, which is further supported by deep profiling of whole proteome and detergent-insoluble proteome in the brain.

MDK is a heparin-binding growth factor of ∼15 kDa, conserved between mouse and human, with 87% protein sequence identity^30^. MDK consists of a signal peptide, an N-terminal domain involved in receptor binding, and a C-terminal domain containing the heparin-binding site. Its structure is stabilized by five intra-domain disulfide bonds, three in the N-terminal domain and two in the C-terminal domain. We expressed recombinant His-tagged MDK proteins in mammalian 293 cells and *E. coli*, and removed the tag by proteases after purification to avoid functional interference (**Extended Data Fig. 1a**). Our top-down and bottom-up MS analysis confirmed the protein sequences and revealed that the MDK expressed in mammalian 293 cells contained all five expected disulfide bonds, while the *E. coli*-expressed version existed as a mixture of three forms, with the five-disulfide form being the most prevalent. The *E. coli*-expressed version showed slightly lower activity in the ThT assay, potentially due to reduced stability in the other two forms lacking one or two disulfide bonds. However, its overall activity remained comparable to that of the 293-expressed MDK. CD and NMR analyses further confirm the inhibitory effect of *E. coli*- expressed MDK on Aβ assembly.

Our results suggest that MDK targets the processes of elongation and secondary nucleation, which are specific pathways during Aβ aggregation. The Aβ aggregation kinetics starting from monomers is inherently complex, including a lag phase, during which a small but critical concentration of amyloid oligomers accumulates, followed by an elongation phase, leading to fibril formation. Additionally, a secondary nucleation pathway occurs on the surface of existing fibrils, promoting further fibril formation^58,59^. Using the AmyloFit interface^35^, we globally fitted our kinetic data to different models (primary nucleation, secondary nucleation, and fragmentation), with secondary nucleation providing the best results. MDK may inhibit Aβ fibrillation similar to sTREM2, which selectively binds to Aβ and primarily suppresses secondary nucleation, with minimal impact on primary nucleation^60^. sTREM2 interacts with Aβ fibrils via two distinct structural surfaces to inhibit secondary nucleation, and similarly, MDK may recognize Aβ through specific interaction sites, potentially modulating fibril formation via comparable mechanisms. Although the exact Aβ- MDK binding interface remains unknown, *in-silico* modeling by AF2 multimer predicts that MDK interacts with Aβ via its C-terminal heparin-binding domain.

Targeting Aβ aggregation is a key therapeutic strategy to slow disease progression. The success of Aβ antibody therapy^14,15^ underscores the importance of Aβ assembly, particularly protofibrils, in AD pathogenesis and highlights the need for precise Aβ conformation targeting to improve therapeutic outcomes^61^. Further research is needed to examine the Aβ-MDK interaction at an atomic level, which could provide critical mechanistic insights for the rational design of effective therapeutic strategies.

In addition to its role in inhibiting Aβ assembly, MDK functions as a growth factor that promote neurite outgrowth and cell survival^30^. It has been shown to interact with several membrane receptors, including PTPζ^62^, LRP-1^63^, integrins (α6β1 and α4β1)^64^, Neuroglycan C^65^, ALK^66^, and Notch-2^67^, regulating development, cell migration and inflammation during brain injury^30^. Although most of these MDK receptors are underexplored in AD, LRP-1 is a receptor for the risk factor *APOE* in AD development^68^. In our proteome analysis of *Mdk* KO mice, we did not observe significant changes in the protein levels of these known MDK receptors. However, we identified alterations in a subset of proteins (WPC3 and WPC4) resulting from the knockout of the *Mdk* gene, independent of the FAD genetic background. These perturbed proteins may also influence AD-related pathology in the knockout mice.

When knocking out the *Mdk* gene in FAD mice, we found a noticeable increase in its paralog protein Ptn in the mouse brain. This upregulation may be driven by at least two possible mechanisms: (i) *Mdk* knockout induces Ptn upregulation at the transcriptional level, (ii) *Mdk* knockout leads to Aβ accumulation, which in turn slows the turnover of Aβ-associated proteins, including Ptn. The first mechanism may not apply to *Mdk* knockout mice without amyloid pathology, as the Ptn protein level remains unchanged under these conditions (**Fig. 6h**). Interestingly, the second mechanism was recently suggested in a proteome turnover study, which explained some RNA-protein inconsistencies in FAD mice^24^, highlighting delayed protein turnover within the amyloid microenvironment. To fully elucidate the relationship between Mdk and Ptn and their contribution to AD pathology, we plan to further investigate these mechanisms using double-knockout mouse models.

During the revision of this manuscript, Levites et al. published a proteomics study on human AD cases, complemented by systematic IHC validations and some functional studies^69^. Their study focused on a conserved network termed the “Aβ amyloid responsome,” which includes MDK and its paralog PTN. They reported that commercially available MDK and PTN proteins promoted fibrillation in a ThT assay, and virus-mediated overexpression of MDK and PTN enhanced parenchymal Aβ amyloid deposition and CAA in CRND8 mice, a transgenic model overexpressing mutant human APP^70^. In contrast, using fully characterized MDK protein from multiple sources, we clearly demonstrate its negative impact on Aβ assembly through an array of detailed biophysical, EM imaging, and NMR studies. The discrepancy in mouse studies may result from differences in models, perturbations, and other experimental variables. Despite these differences, both studies suggest that MDK modulates amyloid accumulation, underscoring the need for further research to reconcile these findings.

Taken together, our findings demonstrate that MDK disrupts Aβ fibril assembly, and its genetic deletion exacerbates plaque accumulation in the FAD mouse model. Given the extended prodromal period before clinical AD onset, early Aβ accumulation may activate compensatory mechanisms to mitigate Aβ toxicity. Our research suggests that MDK may serve as one such factor, providing protective effects against Aβ-induced toxicity and contributing to resistance against amyloid pathology. Further exploration of the MDK pathway could offer new insights into modulating AD disease progression.

## Acknowledgments

We thank Dr. Ines Chen for critical readings and comments. We also thank St. Jude Shared Resources and Core Facilities, including Protein Technology Center, Biomolecular NMR Center, Cryo-electron Microscopy and Tomography Center, Cell and Tissue Imaging Center, Animal Research Center, Center for Advanced Genome Engineering, and Center for Proteomics and Metabolomics. This work was partially supported by the National Institutes of Health grants R01AG053987, RF1AG064909, RF1AG068581, U19AG069701, P30CA021765 and the ALSAC foundation. The Banner Sun Health Research Institute Brain and Body Donation Program was supported by the National Institutes of Health grants U24NS072026, P30AG019610, P30AG072980, the Arizona Department of Health Services, the Arizona Biomedical Research Commission and the Michael J. Fox Foundation for Parkinson’s Research.

## Contributions

J.P., P.-C.C., G.Y. M.Z. conceived this project. M.Z., Y.Y, J.M.Y., Y.H., A.S.T., R.K., C.R.G., M.T., and S.C. engaged in biophysical experiments. S.Y., Z.Wang, K.E.H., H.S., A.H., S.H., Y.J., and P.-C.C. performed biological experiments. P.-C.C., S.Y., Y.J., S.M.P.-M., and V.S. generated and bred the mouse models. S.Y., Z.Wang, D.L., Y, H., Z.Wu, A.A.H., X.W. contributed to the mass spectrometry-based proteomics analysis. G.E.S. and T.G.B. characterized and provided human brain samples. M.Z., Y.H., P.-C.C. and J.P. wrote the manuscript.

## Competing interests

The authors declare no competing interests.

## Methods

All experiments performed in this study comply with all relevant ethical regulations. All animal experiments were approved by the Institutional Animal Care and Use Committee (IACUC) at St Jude Children’s Research Hospital.

### Statistics and reproducibility

Sample sizes were not predetermined using statistical methods; however, they closely resembled those reported in previous publications ^68–70^. While the assumption of normal data distribution was made, formal testing for normality was not analyzed. Variances were assessed using the F-test. An unpaired Student’s *t*-test was used for comparisons between two groups with equal variance, while an unpaired Welch’s *t*-test was used for groups with unequal variance. For proteomics data, *p*-values were calculated by moderated *t*-test in the limma package, followed by false discovery rate analysis by the Benjamini-Hochberg procedure. Animals were utilized whenever available without randomization. Outliers were removed using the Dixon’s Q-test. The investigators were blinded to genotype information during the quantitation of immunostaining for plaque and microglia.

### Human brain samples

Human postmortem brain tissue samples (frontal gyrus) for Western blot analysis were provided by the Brain and Body Donation Program at Banner Sun Health Research Institute. Clinical and pathological diagnoses were based on the established criteria^71^.

### Mouse model

All mice (C57BL/6J) were housed under a 12-h light: 12-h dark cycle at 22-25 °C and 40%-60% humidity. *Mdk* knockout mice were generated using CRISPR-Cas9 technology to partially delete coding region in the genome to cause frame shift mutation. Briefly, dual gRNAs were designed with *in silico* off-target analysis to determine highly unique spacer sequence with at least 3 bp of mismatch to any other site in the mouse genome. In addition, we also consider the targeted region covered among all isoforms, and select CAGE257.MDK.g2 (5’-UGCGGCAUGGGCUUCCGCGA- 3’) and g14 (5’-GCACCUUGCAAUGGACGCGC-3’). Precomplexed ribonucleic proteins (RNPs) consisting of 25 ng/µl of each sgRNA (Synthego) and 50 ng/ µl SpCas9 mRNA (Trilink) were injected into the cytoplasm of C57BL/6J fertilized zygotes and transferred to CD-1 pseudo-pregnant fosters. Resulting animals were genotyped by targeted next generation sequencing using primers CAGE257.F: 5’- GTGAGGCAGGCCGTGTGACCAAGTG-3’ and CAGE257.R:5’-TGCAGTCGGCTGATGGGAGAGTGGC-3’ and analyzed using CRIS.py^72,73^. Founder animals with out-of-frame mutations were backcrossed to C57BL/6J mice for two generations. A founder carrying a 23 bp deletion in exon 3 was selected for the study. Genotyping of the knockout mice was carried out by PCR with primers specific for WT (F: 5’-CTTCCGCGAGGGTACCTG -3’, R: 5’-TATTCGCTGACCTTTGGTCTT -3’) and the *Mdk* knockout (F: 5’- GCACGCCATGTCCTTAACTTT-3’, R: 5’-TGGGTCTGGGCCCATG-3’). FAD mice (B6.Cg-Tg(APPSwFlLon,PSEN1*M146L*L286V)6799Vas/Mmjax) were obtained from Jackson Laboratory (MMRRC_034848-JAX) and underwent backcrossing with C57BL/6J for at least 10 generations. *FAD/Mdk^-/-^* (FAD/KO) mice were generated by crossing with *FAD/Mdk^+/-^* with *Mdk^+/-^* mice. In mouse experiments, similar numbers of male and female animals were utilized.

### Western blotting

Protein lysates underwent separation via 4-20% Tris-Glycine gel and transfer to 0.22 µm nitrocellulose membranes. Blots underwent a 1 h blocking with 3% BSA for MDK or 5% skim milk for other targets. The following primary antibodies were incubated overnight at 4°C: anti-mouse MDK (1:1000, sheep polyclonal, R&D, AF-7769), anti-Aβ (1:1000, mouse monoclonal, clone 82E1, IBL, 10323), and anti-β-tubulin (1:5000, rat monoclonal, clone YOL1/34, abcam, ab6161). After three 5-min washes, blots were exposed to secondary antibodies conjugated with HRP (1:40,000, Jackson ImmunoResearch) for 1 h at 21 °C, followed by another three 5-min washes. Chemiluminescence was developed by using either the SuperSignal West Pico PLUS or Femto substrate (Thermo Fisher Scientific) and detected by ChemiDoc system (Bio-Rad).

Recombinant MDK expression and purification from *E. coli*

The MDK construct (22-143 aa) was overexpressed in BL21 (DE3) Rosetta 2 cells. A single colony was used to cultivate a starter culture overnight. Subsequently, 10 ml of the starter culture was used to inoculate 1 L of 2XYT medium containing 50 μg/ml kanamycin and 35 μg/ml chloramphenicol. The inoculation took place at 37°C with agitation at 220 rpm until the cell density reached an OD₆₀₀ of approximately 0.8.Protein overexpression was induced by adding 0.5 mM isopropyl β-D-thiogalactopyranoside (IPTG), and the culture was further incubated at 18°C for 16 h. Cells were harvested by centrifugation at 5000 rpm for 15 min at 4°C, and the cell pellet was frozen at −80°C until use. The pellet was resuspended in lysis buffer (20 mM HEPES, pH 7.4, 500 mM NaCl, 5% glycerol) supplemented with an EDTA-free complete protease inhibitor cocktail tablet (Roche).

Cell disruption was performed using a cell disruptor (Constant Systems Ltd.) at 20.0 kPSI pressure in two rounds of lysis. The resulting lysate was cleared by centrifugation at 20,000 rpm for 30 min at 4°C and subsequently applied to a gravity column following a 30-min incubation with pre-equilibrated Ni-NTA resin at 4°C. The column was washed with 10 column volumes (CV) of wash buffer A (20 mM HEPES, pH 7.4, 500 mM NaCl, 12.5 mM imidazole, and 5% glycerol), followed by 10 CV of wash buffer B (20 mM HEPES, pH 7.4, 500 mM NaCl, 50 mM imidazole, and 5% glycerol). The protein was eluted with buffer containing 20 mM HEPES, pH 7.4, 500 mM NaCl, 250 mM imidazole, and 5% glycerol.

Eluted fractions were analyzed by SDS-PAGE using a 4–20% gel. Subsequently, the elution fractions were combined, and the His-tag was cleaved from the protein using tobacco etch virus (TEV) protease overnight at 4°C. The cleaved protein was further purified by size-exclusion chromatography using a Superdex 75 16/600 column (Cytiva) equilibrated with 50 mM Tris-HCl, pH 7.5, 500 mM NaCl, and 5% glycerol. Finally, the protein fractions were pooled, concentrated to the desired concentration, flash-frozen in liquid nitrogen, and stored at −80°C for subsequent experiments.

### Recombinant MDK expression and purification from mammalian 293 cells

The MDK gene with an Igκ secretion signal and followed by a His tag, was cloned into the pcDNA 3.4 vector. Recombinant protein was expressed in Expi293F cells (Thermo Fisher Scientific, A14527) through transient transfection using the ExpiFectamine™ 293 Transfection Kit. For the transfection, Expi293F cells at a density of 3 x 10^6^ cells/ml were treated with 1 mg of DNA plasmid per liter of cells. The transfected cells were incubated at 37°C in an orbital shaker set to 140 rpm with 5% CO_2_ for ∼24 h. Then the cells were centrifuged at 1000 x g for 15 mins, and the supernatant was collected. This supernatant was applied to pre-equilibrated Ni-NTA resin (Qiagen) and incubated for 1 h at 4°C. The incubated resin was then collected by centrifugation at 1500 rpm for 10 mins at 4°C and washed with 10 column volumes (CV) of wash buffer A (20 mM HEPES, pH 7.4, 500 mM NaCl, 12.5 mM imidazole, and 5% glycerol), followed by 10 CV of wash buffer B (20 mM HEPES, pH 7.4, 500 mM NaCl, 50 mM imidazole, and 5% glycerol). The protein was eluted using a solution of 20 mM HEPES, pH 7.4, 500 mM NaCl, 250 mM imidazole, and 5% glycerol. The eluted fractions were analyzed by loading onto a 4-20% SDS-PAGE gel. The MDK-containing fractions were pooled, and the His tag was cleaved from the protein overnight at 4°C using tobacco etch virus (TEV) protease. The cleaved protein was then further purified by size exclusion chromatography utilizing a Superdex 75 10/300 Increase column (Cytiva), which was pre-equilibrated with 20 mM sodium phosphate, pH 7.2, 500 mM NaCl, and 5% glycerol. Finally, the protein fractions collected from the column were pooled, concentrated, flash-frozen in liquid nitrogen, and stored at −80°C for subsequent experiments.

### MS analysis of recombinant MDK and Ub proteins

Purified MDK (350 µg/mL, 50 mM Tris-HCl, pH 8.0) was alkylated with 2 mM iodoacetamide for 1 h at RT in the dark to block any potential free thiols. The proteins were then desalted using C18 StageTips (Thermo Scientific) and dried before being re-dissolved in 5% FA for LC-MS analysis. Ubiquitin was prepared in 5% FA directly. LC-MS acquisition was performed using an Ultimate 3000 RSLCnano System coupled to an Q Exactive HF Orbitrap MS (Thermo Fisher Scientific). Samples were separated on a C18 column (75 μm x 20 cm) packed with 1.9 μm C18 resin (Dr. Maisch GmbH). Mobile phase A consisted of 0.2% formic acid, 5% DMSO; Mobile phase B consisted of solvent A plus 65% ACN. The gradient program varied from 20 to 30 min for different proteins. MS data were acquired with a full MS scan with m/z range of 410-1200 or 410-2000 for different proteins, all at R = 240000 (m/z = 200). The resulting raw data were deconvoluted using the Thermo BioPharma Finder Software to obtain the isotopic mass of the intact protein.

For bottom-up analysis, the same samples were alkylated with 2 mM iodoacetamide for 1 h at RT in dark. Subsequently, the proteins were digested in two rounds with trypsin at an enzyme-to-protein ratio of 1:10 (w/w) at 21 °C for at least 8 h. The resulting peptides were desalted using C18 Stage Tips (Thermo Fisher Scientific). LC-MS/MS acquisition was performed using an Ultimate 3000 RSLCnano System coupled to a Q Exactive HF Orbitrap MS (Thermo Fisher Scientific). Samples were redissolved in 5% FA and loaded onto a C18 column (75 μm x 20 cm) packed with 1.9 μm C18 resin (Dr. Maisch GmbH). Mobile phase A consisted of 0.2% formic acid and 5% DMSO, while mobile phase B consisted of solvent A plus 65% ACN. LC separation was performed with a 5-50% buffer B gradient in 25 min. MS analysis was operated in data-dependent acquisition mode with a full MS scan between m/z 410-1200 at R = 60000 (m/z = 200), followed by 10 MS/MS scans at R = 60000 (m/z = 200) with an isolation width of 1.0 m/z. The AGC target for MS and MS/MS scan were set to 3 x 10^6^ and 1 x 10^6^, respectively, with maximum injection time of 50 ms and 110 ms. The precursors with charge states of 2 to 7 were selected for HCD fragmentation with normalized collision energy set to 30%. Dynamic exclusion was set to 10 sec. All samples were analyzed in technical triplicates.

For the identification of disulfide bonds, the raw data were analyzed using pLink 2.0 software (version 2.3.11)^74^ with the following parameters: Cross-linker, SS; Database, FASTA file constructed based on the MDK sequence; Precursor tolerance, 10 ppm; Fragment tolerance, 10 ppm; Number of missed cleavages, 0; Peptide mass, [200, 6000]; Peptide length, [2, 60]; Fixed modification, carbamidomethylation for cysteines; Filter tolerance, ± 10 ppm; FDR, 1% at PSM level. Specifically, the identification of Cys17 and Cys25 containing cross-linked peptides were manually checked, as peptide sequence between Cys17 and Cys25 cannot be cleaved by trypsin. This may allow for the formation of two disulfide bonds among 3 peptides, such as the EGTC_41_GAQTQR–GGPGSEC_17_AEWAWGPC_25_TPSSK–C_50_R, which could not be identified by pLink2. Peptide quantification was performed using the peak area of precursors with the highest intensity. Data were further normalized using the intensity of one trypsin peptide “VATVSLPR”. The top-five most abundant disulfide bonds were shown in figures.

### ThT fluorescence assay

Aβ40 and Aβ42 (rPeptide, #A-1153-1, and #A-1163-2) with a purity of > 97%. Each peptide vial was suspended in 1,1,1,3,3,3-hexafluoro-2-propanol (HFIP), incubated for an h to disperse any preexisting aggregates. Further, these samples were aliquoted and carefully dried in a nitrogen stream and stored in −80°C. Aliquots of purified Aβ species were dissolved in 10 mM NaOH to a concentration of 2 mg/ml followed by 10 min cooling and sonication in an ice water bath for 1 min. The concentration of the final monomeric sample was quantified by nanodrop at 280 nm. MDK protein purified from *E. coli* and 293 cells was dialyzed overnight in 50 mM Tris-HCl buffer, pH 7.5 and final concentration was determined by absorbance measurements at 280 nm. Further, the freshly dissolved Aβ peptides were diluted to 5 µM in absence and presence of different concentrations of purified MDK containing 50 mM Tris-HCl, pH 7.5 buffer containing 20 µM Thioflavin. 100 µl of solution was added into 96-well half area, solid bottom, clear, sterile, microtiter plates and sealed with sealing tape to prevent evaporation. All kinetic experiments were performed at 37°C under quotient conditions at every 5 min in a Clariostar plate reader (BMG Labtech) using an excitation and emission wavelengths of 440 and 480 nm respectively. All these assays were performed in three replicates. The Aβ peptides were diluted to 5 µM at a final concentration in absence and presence of human Ub protein (5 and 10 µM, Bio-Techne R & D systems, U-100H-10M) to monitor aggregation kinetics as described above. Aggregation kinetics of MDK and Ub at different concentrations in the same buffer (50 mM Tris-HCl) with fixed ThT concentrations (20 µM) were also recorded.

### Circular dichroism spectroscopy

All CD spectra were recorded on a JASCO-1500 Spectrophotometer at a wavelength of 195 - 250 nm with a step size of 0.1 nm, 1 nm bandwidth, and a scan speed of 50 nm/min. Aβ40/42 samples with a final monomeric concentration of 10 µM were incubated in the absence and presence of MDK (1:2) in a similar buffer. The samples were incubated at 37 °C and studied in a time course (up to 24 h), where a stable plateau value was reached. A spectrum of buffer was recorded separately and subtracted from the spectra of Aβ40/42. Spectra of MDK protein were recorded separately and subtracted from the spectra of Aβ and MDK, and the CD signal was given as ellipticity. Aβ (10 µM) samples incubated in absence and presence of ubiquitin (1:2) were also recorded under identical conditions. Moreover, CD spectra of MDK and ubiquitin were independently recorded at different time intervals and subtracted from their respective buffers. For each sample, the average of five scans was recorded and subsequently smoothed using Savitzky-Golay smoothing method and reported.

### Negative stain transmission electron microscopy

Aβ species alone and reactions involving Aβ together with 10 µM of MDK samples from the kinetic experiments were collected at the endpoint of the ThT experiments. Additionally, Aβ samples, both in the absence and presence of 10 µM of ubiquitin, were collected at the endpoint of the ThT experiments. 400-mesh copper grids (CF400-Cu grids, Electron Microscopy Sciences) were plasma cleaned with an Ar/O2 gas mixture for 10s using Solarus plasma cleaner (Gatan), followed by 5 µl of sample. The samples were allowed to absorb for 1 min before blotting away the excess liquid, followed by rinsing with Milli Q water and subsequent staining using three successive applications of 2% uranyl acetate. The last round of stain application was allowed to sit for 1 min before blotting away the excess stain. The grids were air-dried prior to imaging using a 120 kV Talos L120C TEM (Thermo Fisher Scientific) equipped with a CETA detector (TFS).

### Immunogold negative-stain electron microscopy

Extraction of Aβ filaments was performed as in the previous report^75^. Aβ filaments were deposited on glow-discharged 400 mesh formvar/carbon coated copper grids (EM Sciences CF400-Cu) for 40 s. Subsequently, the grids were blocked for 10 min with 1% BSA in PBS and incubated with anti-MDK (1:50, goat polyclonal, Bio-Techne, #AF-258-SP). After rinsing with blocking buffer, the grids were incubated with anti-goat IgG conjugated with 10 nm gold particle (1:20, Sigma), followed with wash and stained with 2% uranyl acetate for 1 min. Images were acquired at 11,000x with a Gatan Orius SC200B CCD detector on a Tecnai G2 Spirit at 120 kV.

### NMR HSQC Spectroscopy

The ^15^N-labeled Aβ40 and Aβ42 peptides (rPeptide, #A-1101-2 and #A-1102-2) were dissolved in 10 mM NaOH to a concentration of 2 mg/ml, followed by 10 min of cooling and 1 min of sonication in an ice water bath. After aliquoting on ice, the samples were stored in a −80°C freezer. Subsequently, 10 µM of the ^15^N-labeled Aβ42 and Aβ40 peptides were incubated with 0, 5, and 10 µM MDK in low-binding tubes (Eppendorf, #0030108434) at 37°C for 24 and 48 h, respectively. In addition, for control experiments Aβ42 and Aβ40, samples were incubated independently in the absence and presence of ubiquitin (10 µM) at 37°C for 24 and 48 h, respectively. The NMR spectra of all these samples including 10 µM monomeric Aβ species, were acquired on Bruker Avance 600 MHz or 800 MHz at 278K. The spectrometers are equipped with a triple-resonance cryoprobe, and the data were processed in NMRPipe^76^. and analyzed using NMRFAM-SPARKY^77^. The backbone resonance assignment of Aβ40 was taken from BMRB (ID: 17795). Binding of MDK to the 15N-labeled Aβ40 and Aβ42 was investigated using two-dimensional (2D) [^1^H-^15^N] so fast HMQC^78^ spectrum recorded either with 32 scans or 128 scans with an interscan delay of 0.2 s. The intensity of the peaks from different spectra was quantified, normalized with respect to the highest peak and used in analyzing the data.

### Preparation of Sarkosyl-soluble and -insoluble proteins from mouse brain

The analysis procedure was adapted from a previously published method^13^. Mouse cortices were lysed in a low-salt buffer (50 mM HEPES, pH 7.4, 1× protease inhibitors (Thermo Fisher Scientific), 10:1 (v/v) ratio), and then mixed with a Sarkosyl-containing buffer (10-fold dilution) for differential extraction. For ELISA analysis, a stringent buffer was used to enrich for Aβ fibrils (50 mM HEPES, pH 7.4, 100 mM NaCl, 5 mM EDTA, 1.0% Triton X-100, 0.5% sodium deoxycholate, 1% Sarkosyl, 10% glycerol, 1× protease inhibitors). For proteomics analysis, Sarkosyl was reduced to 0.2% to retain the proteins associated with Aβ fibrils. Lysates were centrifuged at 3,000 × g for 5 min to remove cell debris, and the supernatant was subjected to ultracentrifugation at 55,000 rpm for 1 h at 4°C. The final supernatant was collected as Sarkosyl-soluble fraction, and the pellet was collected and solubilized by a urea-containing buffer (50 mM HEPES, pH 8.5, 8 M urea, and 0.5% sodium deoxycholate) as Sarkosyl-insoluble fraction.

### ELISA analysis

Aβ levels in the Sarkosyl-soluble and insoluble fractions of mouse brains were measured using human Aβ40 and Aβ42 ELISA kits (Invitrogen, KHB3481 and KHB3441). Briefly, the strips in a 96-well plate were pre-coated with a mouse monoclonal Ab specific to the N-terminus of human Aβ. Diluted protein samples (∼200-fold Sarkosyl-soluble samples, and ∼20,000-fold Sarkosyl-insoluble samples) were added into the plate and incubated overnight at 4°C together with another Ab specific to the C-terminus of human Aβ40 or Aβ42. The bound primary antibody was then detected using an HRP-conjugated secondary antibody, followed by a 1 h incubation at 21 °C. After washing, a chromogenic substrate solution was added and OD_450_ was measured using a microplate reader (Epoch, BioTek).

### Immunohistochemistry

The staining was modified from previously report^12^. Brain tissue samples were fixed with 10% formalin and embedded in paraffin. 10 µm sagittal sections were deparaffinized, rehydrated and rinsed with water. Antigen retrieval was performed with 10 mM citric buffer (pH 6.0) with boiled water bath for 20 min and cooled down to 21 °C. Endogenous peroxidase activity is further blocked by 1% H_2_O_2_ in PBS buffer for 10 min. After rinsed by PBS, sections were then blocked with 5% donkey serum in PBS with 0.3% Triton X-100 (PBST) for 30 min at 21 °C followed with primary antibodies diluted with PBST plus 2% BSA and 0.2% skim milk for overnight at 4°C: anti-human MDK, (goat polyclonal, 1:200, R&D, AF-258-SP), anti-mouse MDK (1:200, sheep polyclonal, R&D, AF-7769), anti-Aβ (1:100, clone 82E1, IBL, 10323), anti-IBA1 (1:400, rabbit polyclonal, Fujifilm Wako, 019-19741). After washing with PBS, sections were then incubated with secondary antibodies (1:500, Jackson ImmunoResearch) for 1 h at 21 °C. For MDK, we used HRP polymer-conjugated secondary antibodies (Vector Laboratories, MP- 7405), followed with signal amplification using TSA-Cy5 (1:200, Akoya Biosciences).

For fibrillar plaque staining, X34 dye was used due to its high specificity for binding to β-sheet structures. After antigen retrieval, slides were stained with 1 mM X34 dye for 10 min, followed by washes with 40% ethanol and PBS. For chromogenic immunodetection, biotin-conjugated secondary antibodies were used followed with ABC reaction by Vectastain ABC kit (Vector Laboratories, PK-4000) and developed by 3,3-diaminobenzidine solution (Vector Laboratories, SK-4100). Images were captured by Zeiss Axioscan.Z1 and Zeiss LSM 780 confocal microscopy.

### Quantitation of Aβ plaques and microglia

Brain tissues were serially sectioned sagittally at 10 µm thickness. Three slides, each representing every 30^th^ section, were utilized for immunohistochemistry followed with imaging captured with Zeiss Axioscan.Z1. QuPath, an open-source software^79^ was carried out for automatic quantitation of density and area of Aβ plaques and microglia in an investigator blinded manner.

### Mass spectrometry-based proteomics

We used an optimized protocol of TMT-LC/LC-MS/MS for deep proteome profiling^48,49^. Protein samples were lysed by homogenization in the lysis buffer (50 mM HEPES, pH 8.5, 8 M urea, and 0.5% sodium deoxycholate), and their concentrations were measured by the BCA assay (Thermo Fisher Scientific, #23227) and confirmed by Coomassie-stained short SDS gels ^80^. Quantified protein samples (∼0.1 mg per TMT channel) were digested with Lys-C (1:100 w/w, Wako, distributor, #121-05063) for 2 h at 21 °C, followed by dilution to decrease urea to 2 M and trypsin digestion (1:50 w/w, Promega, #V5113) overnight at 21 °C. Cys residues were reduced and alkylated by iodoacetamide. The proteolysis was terminated by adding trifluoroacetic acid to 1%. The resulting peptides were desalted with the Sep-Pak C18 cartridges (Waters), TMT-labeled (Thermo Fisher Scientific, #A34808 and A44520), and pooled equally. The pooled peptides were resolved by basic pH reverse phase LC on an XBridge C18 column (3.5 μm beads, 4.6 mm x 25 cm, Waters; buffer A: 10 mM ammonium formate, pH 8.0; buffer B: 95% acetonitrile, 10 mM ammonium formate, pH 8.0, ∼2 h gradient, 40- concatenated fractions collected)^81^. Each fraction was analyzed by acidic pH LC-MS/MS (75 µm x ∼20 cm, 1.9 µm C18 resin from Dr. Maisch GmbH, buffer A: 0.2% formic acid, 5% DMSO; buffer B: buffer A plus 65% acetonitrile, ∼1.5 h gradient). The settings of Q Exactive HF Orbitrap MS (Thermo Fisher Scientific) included the MS1 scan (∼410-1600 *m/z*, 60,000 resolution, 1 x 10^6^ AGC and 50 ms maximal ion time) and 20 data-dependent MS2 scans (fixed first mass of 120 *m/z*, 60,000 resolution, 1 x 10^5^ AGC, ∼110 ms maximal ion time, HCD, 32-35% normalized collision energy, ∼1.0 *m/z* isolation window with 0.3 *m/z* offset, and ∼15 s dynamic exclusion)^49^. The raw MS data were searched against protein database by the JUMP software (v1.13.4)^82^, which utilizes both pattern matching and *de novo* tag scoring to improve the sensitivity and specificity. A composite target/decoy database was used to evaluate FDR in peptide identification^83,84^. The protein target database combined downloaded Swiss-Prot, TrEMBL, and UCSC databases (human: 83,955 entries; mouse: 59,423 entries). Search parameters included precursor/product ion mass tolerance (± 10 ppm), full trypticity, static mass shift (TMT tags of 229.16293 304.2071453 and Cys carbamidomethylation of 57.02146 on cysteine), dynamic mass shift (Met oxidation of 15.99492), two maximal miscleavage sites, and three maximal modification sites. Peptide-spectrum matches (PSMs) were filtered by matching scores and mass accuracy to limit protein FDR below 1%.

Proteins were quantified from TMT reporter ions based on our published method^50^. Briefly, TMT reporter ion intensities were extracted for each PSM, corrected by isotopic distribution of TMT reagents, filtered to remove poor PSMs (e.g., minimum intensity of 1,000), and adjusted to alleviate sample pooling bias. The relative protein intensities were averaged from all assigned PSMs after removing outliers (e.g., Dixon’s Q-test or generalized extreme Studentized deviate test). Finally, the absolute protein intensities were derived by multiplying the relative intensities by the grand mean of top three abundant PSMs.

### Weighted Gene Co-expression Network Analysis (WGCNA) analysis

The network analysis was performed using WGCNA R package^51^ using differentially expressed proteins. First, a signed Pearson correlation matrix was computed based on the protein abundance data in all mouse samples, and transformed into an adjacency matrix by applying a soft-threshold power of 16, according to modified scale-free topology criterion^85^. Hierarchical clustering was then performed using the dynamic tree-cutting algorithm to detect initial modules (clusters), which were subsequently merged at a threshold of 0.1 (indicating high similarity on a 0 to 1 scale). Proteins were ultimately assigned to the cluster with the highest correlation, with any ambiguities resolved through manual curation.

### Pathway enrichment by Gene Ontology and KEGG databases

Pathway enrichment analysis was carried out by the Cluster profiler (v4.10.0)^86^ to identify the biological functions of dysregulated genes/proteins in a given dataset. In the analysis, we evaluated whether certain biological processes or pathways were statistically overrepresented in the list of input proteins compared to the background set of mouse proteome. The “enrichment” refers to the increased occurrence of specific functional categories within the input list beyond what would be expected by chance. The analysis was performed using Fisher’s exact test against the Gene Ontology (GO) biological process, molecular function, and cellular component annotations, and KEGG pathway database, separately. The *p* values derived from Fisher’s exact test were further adjusted into FDR using the Benjamini-Hochberg procedure for multiple testing. Enriched pathways with FDR < 10% were considered statistically significant.

### Quantitation and statistical analysis

In small-scale analyses, two-tailed unpaired Student’s *t*-test was used for two-sample comparisons (GraphPad Prism 7.0.5). Proteomics analysis for differentially expressed proteins primarily utilized the limma R package (v3.58.1)^87^ in several steps: (i) obtain the protein quantification data from MS as described above, (ii) perform a log transformation of the data^88^, (iii) calculate the *p* values by moderated *t*-test, and FDR values by the Benjamini-Hochberg procedure, using limma R package, (iv) calculate the mean for each protein under different conditions and derive the log2(fold change), (v) fit the log2(fold change) data of all proteins to a Gaussian distribution to generate a ‘global’ standard deviation value. Statistically significant changes were typically determined using an FDR cutoff of 5% and a log2 (fold change) cutoff equivalent to two standard deviations.

## Data availability

The mass spectrometry proteomics data were deposited to the ProteomeXchange Consortium via the PRIDE partner repository with the dataset identifiers PXD046539 and PXD061103 (the whole proteome of AD mouse models), and PXD045746 and PXD061104 (the detergent-insoluble proteome of the mice).

---

**Extended Data Fig. 1.**
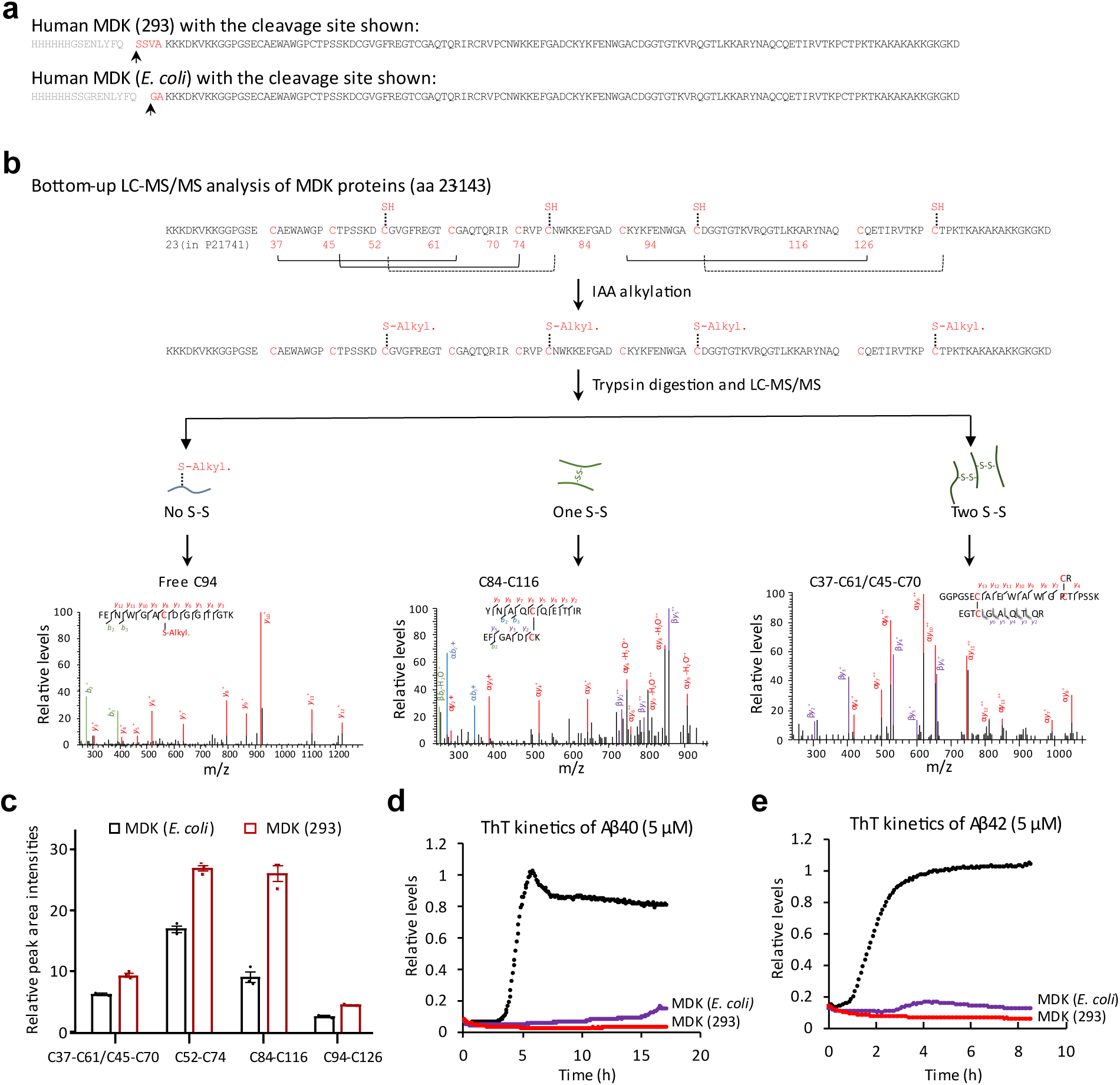
Bottom-up MS analysis of recombinant MDK proteins and their effect on Aβ fibrillation. **a**, Protein sequences of MDK proteins expressed in mammalian 293 or *E. coli* cells. The N-terminal His-tag was cleaved after expression by TEV protease. **b,** Workflow of bottom-up LC-MS/MS analysis to characterize the positions of disulfide (S-S) bonds. Proteins were subjected to IAA alkylation, trypsin digestion and LC-MS/MS analysis. Crosslinked peptides containing single or double S-S bonds were identified. **c,** Relative peak intensities of top five crosslinked peptides (*n* = 3 replicates). **d-e,** Effect of 293- or *E. coli*-expressed MDK (10 µM) on Aβ40/42 (5 µM) fibrillation kinetics in ThT fluorescence assays. Results represent the average of 3 replicates.

**Extended Data Fig. 2.**
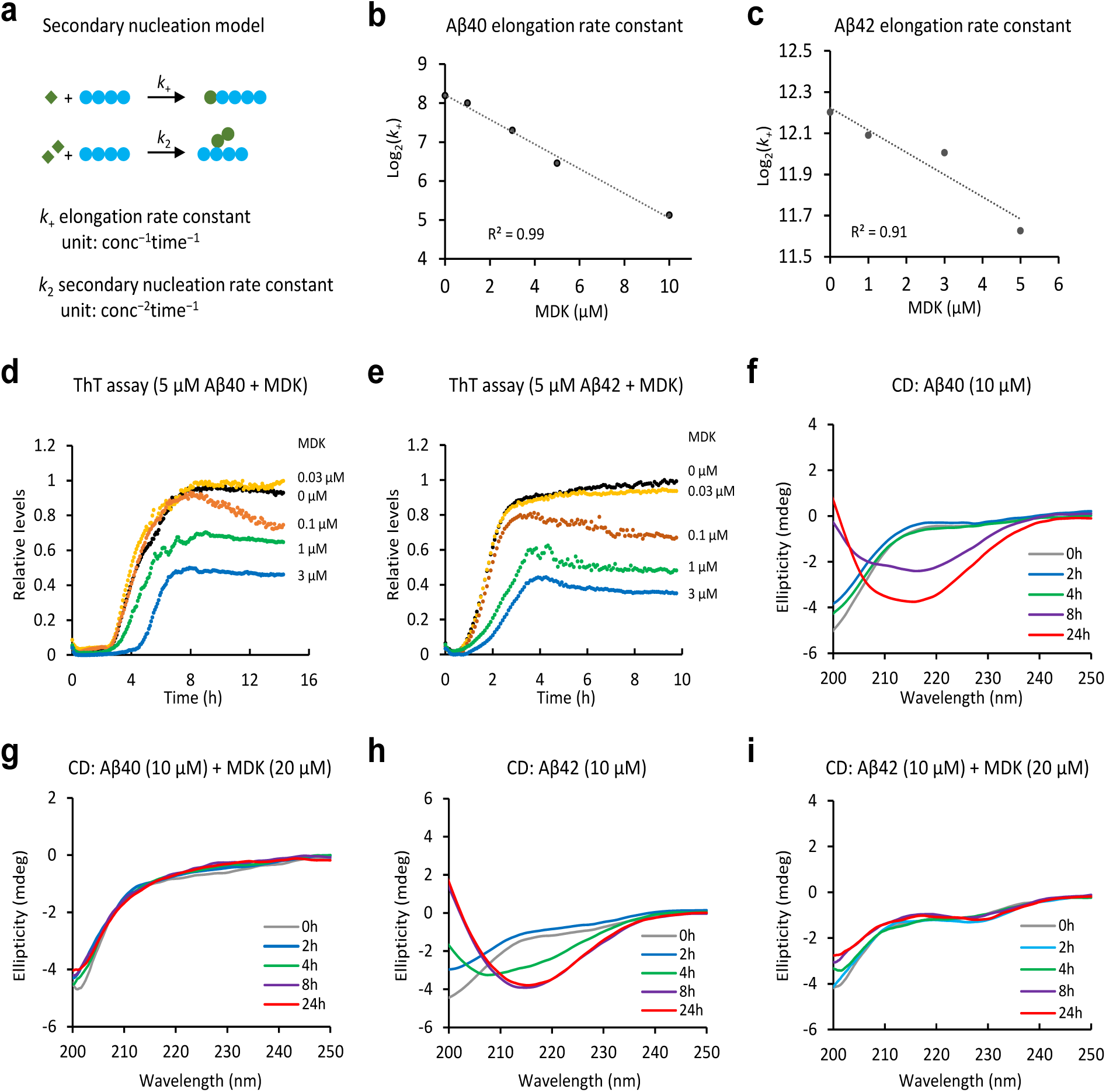
Characterization of the effect of MDK on Aβ fibrillation. **a,** Diagram of the secondary nucleation model illustrating Aβ assembly. The rate constants (*k*_+_ and *k*_2_) are defined with units, reflecting their dependency on protein concentration and time. **b-c,** Analysis of Aβ40 and Aβ42 elongation constants (*k*_+_) by fitting the ThT data. **d**, Effect of MDK (0-3 µM) on Aβ40 fibrillation kinetics (*n* = 3 replicates, averaged data shown). **e,** Effect of MDK (0-3 µM) on Aβ42 fibrillation kinetics (*n* = 3 replicates, averaged data shown). **f,** Time course CD spectra of Aβ40 alone (0-24 h). **g,** Time course CD spectra of Aβ40 with MDK (0-24 h). **h,** Time course CD spectra of Aβ42 alone (0-24 h). **i,** Time course CD spectra of Aβ42 with MDK (0-24 h).

**Extended Data Fig. 3.**
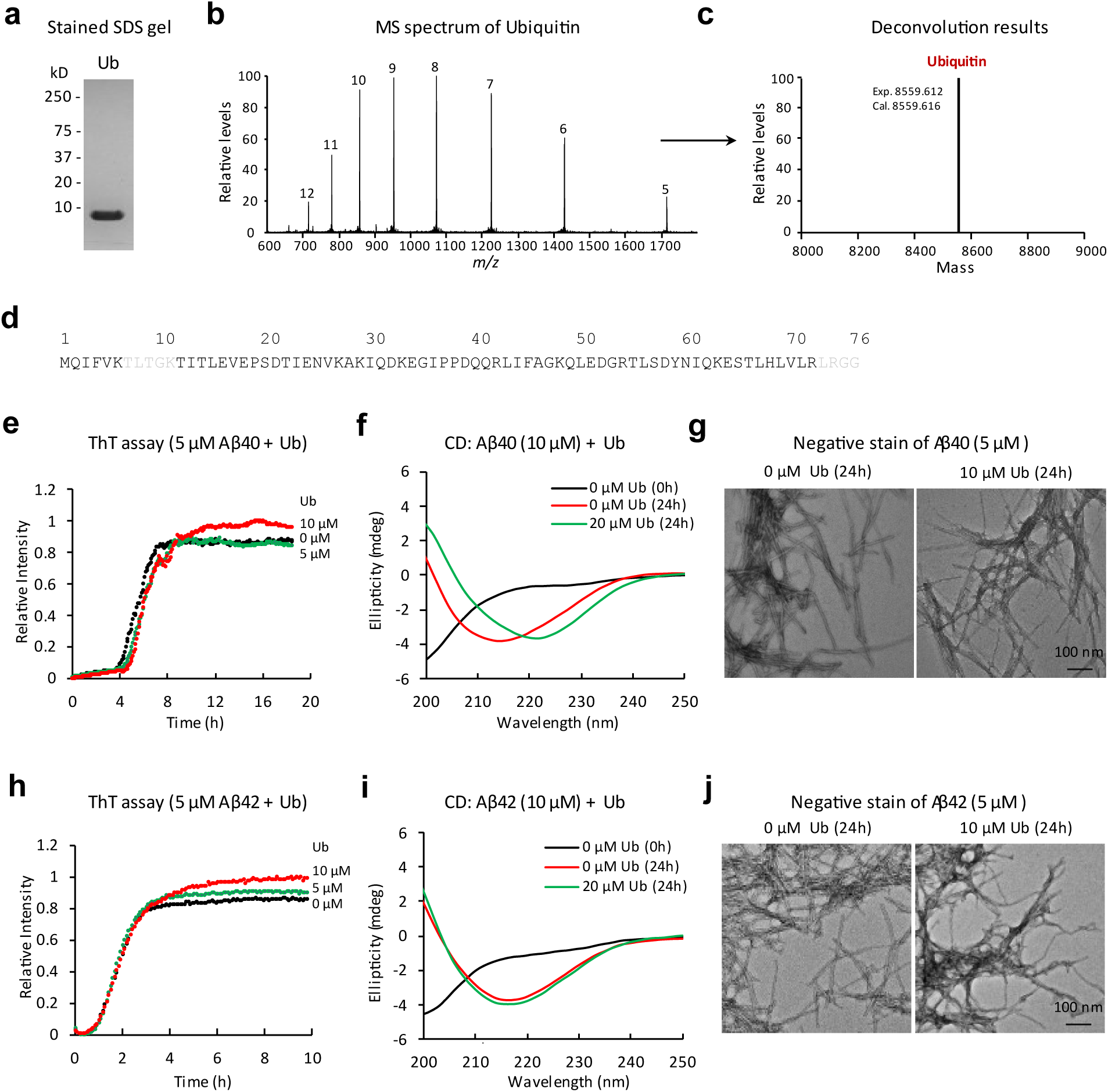
Characterization of ubiquitin protein by MS and its lack of effect on Aβ40/42 fibrillation. **a**, Purified Ub protein on a stained SDS gel. **b-c,** Top-down mass spectrum and deconvoluted results of Ub protein showing different charge states. **d,** Bottom-up LC-MS/MS analysis of Ub, covering the full Ub sequence except two short tryptic peptides (in grey). **e,** ThT fluorescence assay measuring Aβ40 fibril formation at different Ub concentrations (*n* = 3 replicates, averaged data shown). **f**, CD spectroscopy of Aβ40 with or without Ub, with ellipticity reported in millidegrees (mdeg). **g**, Negative stain EM of Aβ40/Ub samples with a scale bar (100 nm). **h**, ThT fluorescence assay measuring Aβ42 fibril formation at different Ub concentrations (*n* = 3 replicates, averaged data shown). **i.** CD spectroscopy of Aβ42 with or without Ub. **j,** Negative stain EM of Aβ42/Ub samples with a scale bar (100 nm).

**Extended Data Fig. 4.**
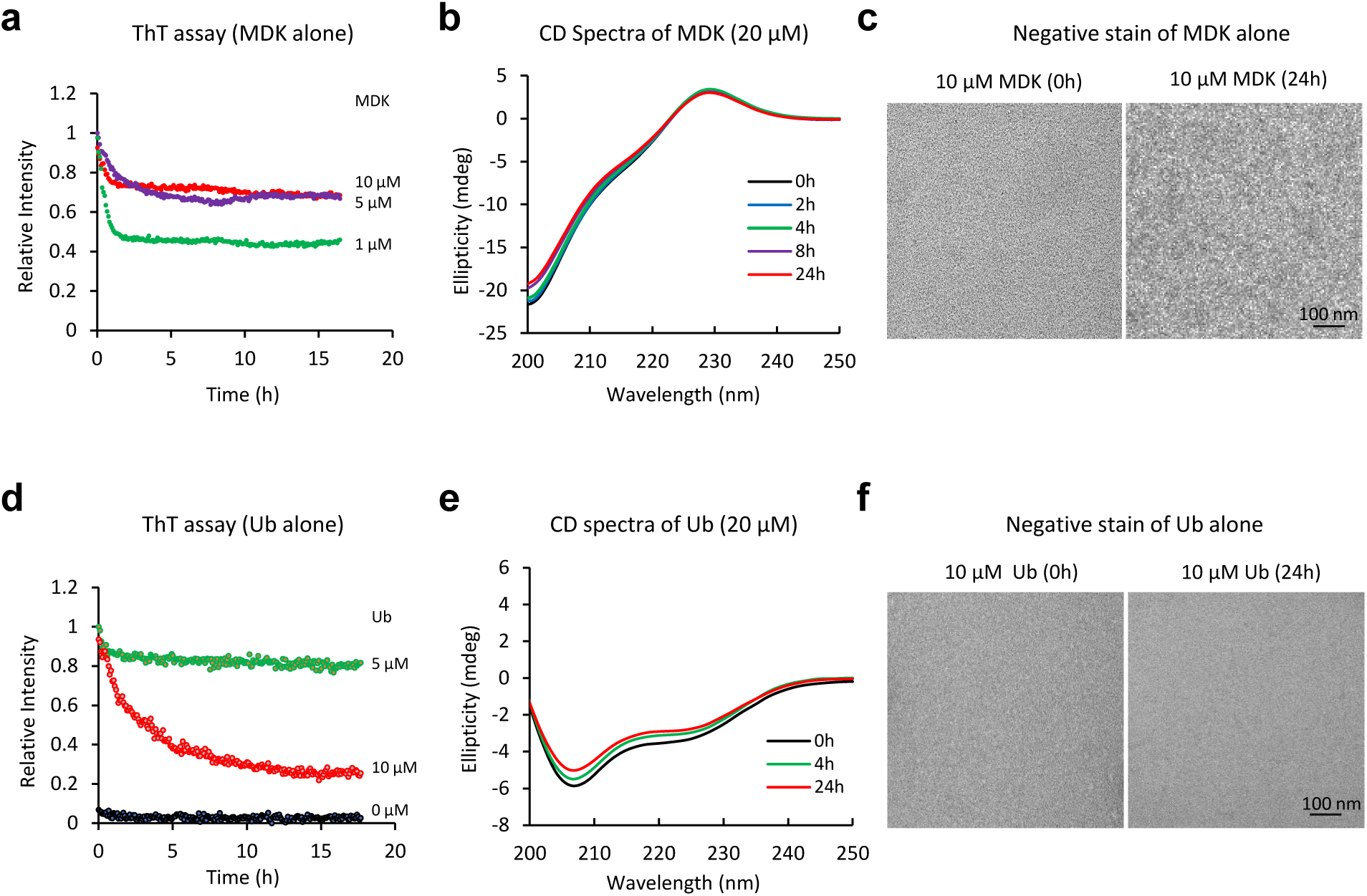
MDK or ubiquitin cannot independently assemble into fibrils. **a**, ThT assay of MDK alone at different concentrations (*n* = 3 replicates, averaged data shown). **b**, Time course CD spectra of MDK (0-24 h). **c**, Negative stain EM of the MDK samples, with a scale bar (100 nm). **d**, ThT assay of Ub alone at different concentrations (*n* = 3 replicates, averaged data shown). **e**, Time course CD spectra of Ub (0-24 h). **f**, Negative stain EM of the Ub samples, with a scale bar (100 nm).

**Extended Data Fig. 5.**
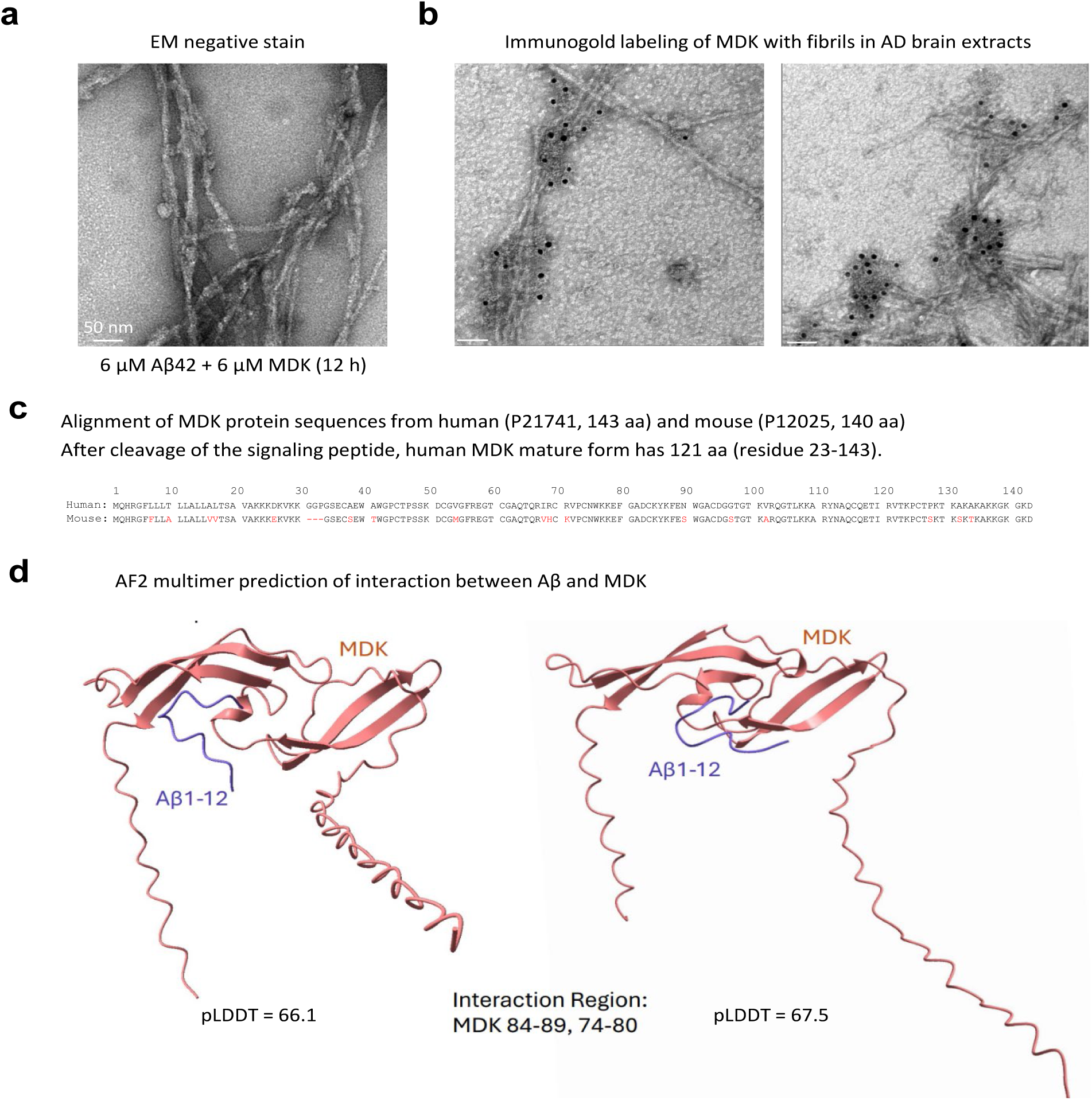
EM images of native Aβ filaments from patients with MDK and structural prediction of Aβ-MDK interaction. **a**, Negative stain EM of the fibril form of Aβ42 (6 µM) with MDK (6 µM), with scale bars (50 nm). **b**, Immunogold labeling of MDK in human brain fibrils. Enriched Aβ filaments from human AD brain lysates were analyzed with anti-MDK goat antibodies, with scale bars (50 nm). **c**, The alignment of human and mouse MDK protein sequences. MDK is a secreted protein containing a signal peptide for secretion (aa 1–22) and the main protein chain (aa 23–143). **d**, AlphaFold2-multimer predicted structures of the Aβ-MDK potential interacting interfaces. The confidence of the predicted 3D structures is evaluated by the predicted Local Distance Difference Test (pLDDT).

**Extended Data Fig. 6.**
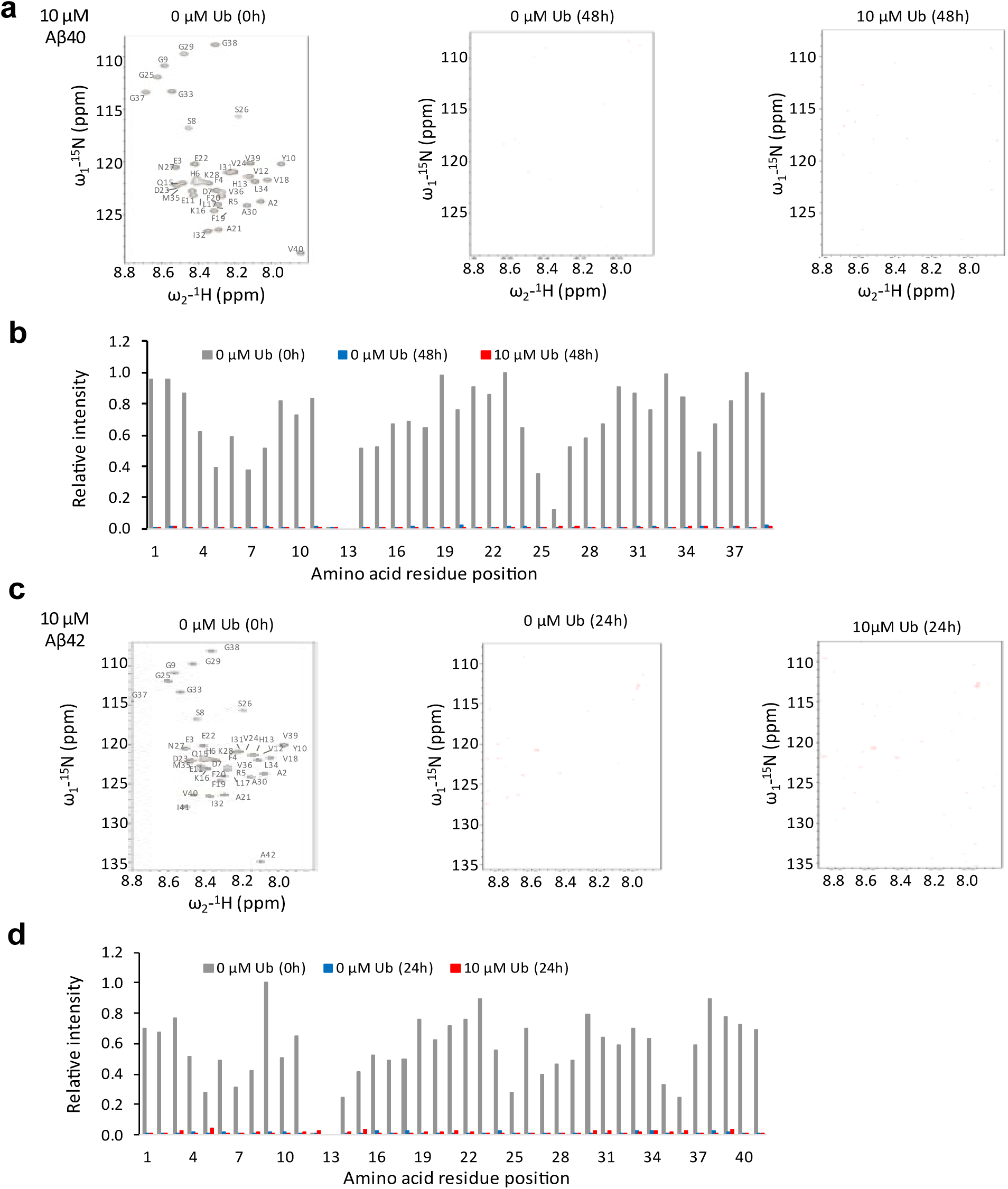
Ubiquitin cannot rescue NMR signals of Aβ peptides. **a,** ^1^H-^15^N HSQC spectra of Aβ40 (10 μM) with or without Ub (10 μM) in 50 mM Tris buffer (pH 7.5). Spectra were collected before incubation (left panel) and after 48 h incubation (right two panels). **b,** Relative cross-peak intensities for each residue, excluding D1 and H14. Intensities were normalized to a maximum value of 1. **c-d,** NMR analysis of Aβ42 under similar conditions as Aβ40, but with 24 h incubation.

**Extended Data Fig. 7.**
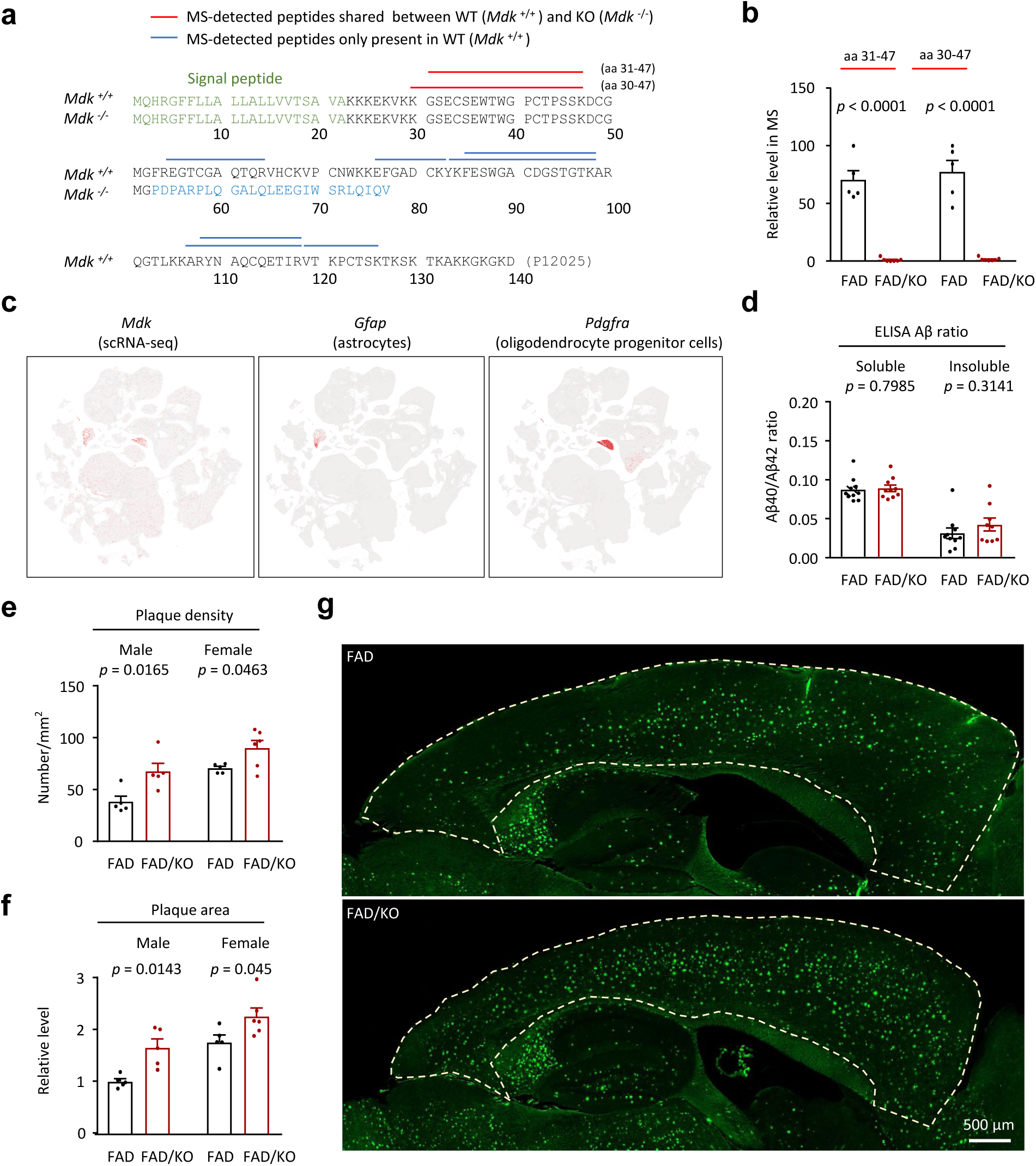
Analysis of MDK expression in KO mice, its cell-specific expression, and its impact on Aβ in male and female mice. **a**, Predicted mouse MDK protein sequences in WT and CRISPR-mediated KO mice, with N-terminal signal peptide shown in green. The 23 base pair deletion in MDK gene alters the open reading frame in the KO mice, potentially generating a truncated protein with distinct C-terminus (in blue) from WT. Peptides detected by MS in brain proteomic analysis are also shown, with shared peptides between MDK and the potential truncated form highlighted in red. The two peptides (aa 30-47 and aa 31-47) allow quantification of MDK full length proteins and the tentative N-terminal truncation. **b**, Quantification of two shared peptides in FAD and FAD/KO. The intensities were normalized by setting the highest value to 100. The absence of their signals in FAD/KO mice strongly suggests that the tentative N-terminal truncation was not expressed at a detectable level. **c**, Expression of *Mdk*, *Gfap* (astrocyte marker) and *Pdgfra* (oligodendrocyte progenitor cell marker) in mouse brain, shown in Uniform Manifold Approximation and Projection (UMAP) plots. The cell-type clusters were derived from single-cell RNA sequencing of more than 1.2 million cells in the whole cortex and hippocampus using the 10x Genomics Chromium platform^44^. Expression levels are indicated by color intensity (red). Mdk is highly expressed in the 377_Astro cluster, with a trimmed mean (25-75%) Log₂(CPM+1) value of 4.13, as well as in the 366_Oligo cluster, with a trimmed mean value of 0.72. **d**, ELISA analysis of Aβ40/Aβ42 ratio in the Sarkosyl-soluble and -insoluble fractions of mouse cortices (*n* = 10 per group). **e**, Quantification of X34-positive amyloid plaque density in the cortices of male (FAD *n* = 5; FAD/KO *n* = 5) and female (FAD *n* = 5, FAD/KO *n* = 6). **f**, Quantification of X34-positive amyloid plaque area in the cortices of the same sets of male and female mice. The results were normalized by setting the mean value of FAD mice to 1. Statistical significance between FAD and FAD/KO in male or female was analyzed by two-tailed Student’s *t*-test. **g**, Example images of X34 staining showing amyloid plaque quantification in the cortices of FAD and FAD/KO mice. Cortices were outlined by the white lines. Scale bar, 500 µm. Data are shown as mean ± SEM in panels **b**, **d**, **e**, **f**. Full statistical information is available in Source Data.

**Extended Data Fig. 8.**
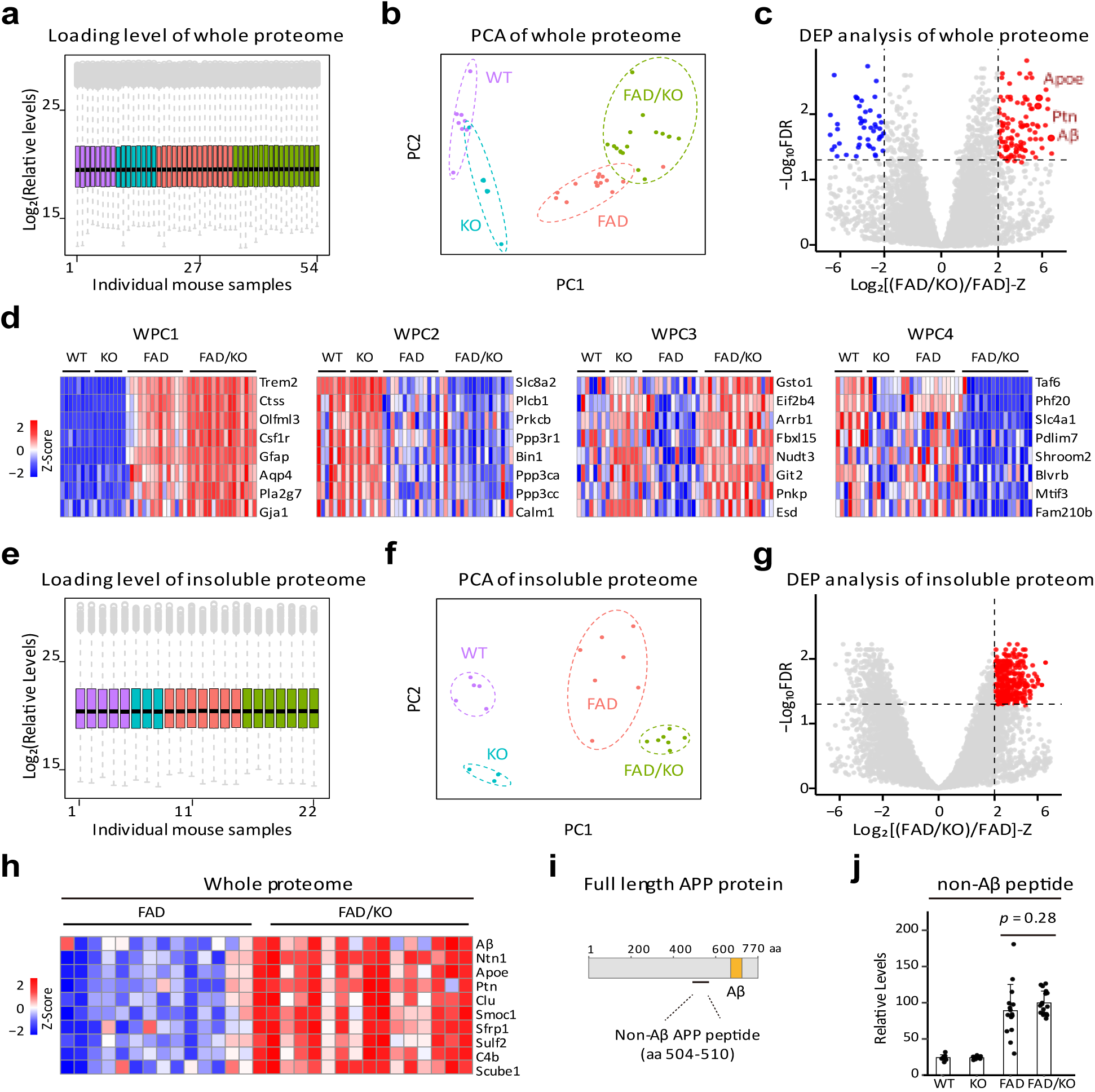
Proteomic comparison of four genotypes derived from FAD and *Mdk* KO mice. **a**, Boxplot of whole proteome data showing equal loading across samples, with WT, KO, FAD and FAD/KO represented in purple, blue, orange, and green, respectively. **b**, PCA of whole proteome data using DEPs, showing genotype-specific separation. **c**, Representative volcano plot showing DEPs between FAD and FAD/KO mice, with FDR values from limma analysis and log2 ratios converted to z values. **d**, Selected DEPs in four whole proteome clusters (WPC), with each protein represented by a colored box after log2 conversion and Z score analysis. **e**, Boxplot of insoluble proteome data. **f**, PCA of insoluble proteome data using DEPs. **g**, Volcano plot comparing the insoluble proteome of FAD and FAD/KO mice. **h,** Heatmap of selected DEPs shared between the insoluble and whole proteomes. **i,** Full-length APP protein diagram showing the position of a representative non-Aβ tryptic peptide. **j,** Quantification of the non-Aβ peptide levels in whole proteome across four genotypes (WT *n* = 8, KO *n* = 8, FAD *n* = 15, FAD/KO *n* = 17), with statistical significance determined by a two-tailed Student’s *t*-test.

## Notes

### Competing Interest Statement

The authors have declared no competing interest.

## References

1 Alzheimer’s_Association. 2023 Alzheimer’s disease facts and figures. Alzheimers Dement 19, 1598–1695, (2023).

2 Bellenguez, C., Grenier-Boley, B. & Lambert, J. C. Genetics of Alzheimer’s disease: where we are, and where we are going. Curr Opin Neurobiol 61, 40–48, (2020).

3 Bellenguez, C. et al. New insights into the genetic etiology of Alzheimer’s disease and related dementias. Nat Genet 54, 412–436, (2022).

4 Wightman, D. P. et al. A genome-wide association study with 1,126,563 individuals identifies new risk loci for Alzheimer’s disease. Nat Genet 53, 1276–1282, (2021).

5 Hardy, J. & Selkoe, D. J. The amyloid hypothesis of Alzheimer’s disease: progress and problems on the road to therapeutics. Science 297, 353–356, (2002).

6 Ballatore, C., Lee, V. M. & Trojanowski, J. Q. Tau-mediated neurodegeneration in Alzheimer’s disease and related disorders. Nat Rev Neurosci 8, 663–672, (2007).

7 Neumann, M. et al. Ubiquitinated TDP-43 in frontotemporal lobar degeneration and amyotrophic lateral sclerosis. Science 314, 130–133, (2006).

8 Nelson, P. T. et al. Limbic-predominant age-related TDP-43 encephalopathy (LATE): consensus working group report. Brain 142, 1503–1527, (2019).

9 Bai, B. et al. U1 small nuclear ribonucleoprotein complex and RNA splicing alterations in Alzheimer’s disease. Proc Natl Acad Sci U S A 110, 16562–16567, (2013).

10 Bai, B. et al. Integrated Approaches for Analyzing U1-70K Cleavage in Alzheimer’s Disease. J Proteome Res 13, 4526–4534, (2014).

11 Hales, C. M. et al. U1 small nuclear ribonucleoproteins (snRNPs) aggregate in Alzheimer’s disease due to autosomal dominant genetic mutations and trisomy 21. Mol Neurodegener 9, 15, (2014).

12 Chen, P. C. et al. Alzheimer’s disease-associated U1 snRNP splicing dysfunction causes neuronal hyperexcitability and cognitive impairment. Nat Aging 2, 923–940, (2022).

13 Zaman, M. et al. Dissecting Detergent-Insoluble Proteome in Alzheimer’s Disease by TMTc-Corrected Quantitative Mass Spectrometry. Mol Cell Proteomics 22, 100608, (2023).

14 van Dyck, C. H. et al. Lecanemab in Early Alzheimer’s Disease. N Engl J Med 388, 9–21, (2023).

15 Sims, J. R. et al. Donanemab in Early Symptomatic Alzheimer Disease: The TRAILBLAZER-ALZ 2 Randomized Clinical Trial. JAMA 330, 512–527, (2023).

16 Bai, B. et al. Deep Multilayer Brain Proteomics Identifies Molecular Networks in Alzheimer’s Disease Progression. Neuron 105, 975–991, (2020).

17 Wang, H. et al. Integrated analysis of ultra-deep proteomes in cortex, cerebrospinal fluid and serum reveals a mitochondrial signature in Alzheimer’s disease. Mol Neurodegener 15, 43, (2020).

18 Higginbotham, L. et al. Integrated proteomics reveals brain-based cerebrospinal fluid biomarkers in asymptomatic and symptomatic Alzheimer’s disease. Sci Adv 6, eaaz9360, (2020).

19 Johnson, E. C. B. et al. Large-scale deep multi-layer analysis of Alzheimer’s disease brain reveals strong proteomic disease-related changes not observed at the RNA level. Nat Neurosci 25, 213–225, (2022).

20 Sathe, G. et al. Quantitative proteomic analysis of the frontal cortex in Alzheimer’s disease. J Neurochem 156, 988–1002, (2021).

21 Roberts, J. A. et al. A brain proteomic signature of incipient Alzheimer’s disease in young APOE epsilon4 carriers identifies novel drug targets. Sci Adv 7, eabi8178, (2021).

22 Bai, B. et al. Proteomic landscape of Alzheimer’s disease: novel insights into pathogenesis and biomarker discovery. Mol Neurodegener 16, 55, (2021).

23 Askenazi, M. et al. Compilation of reported protein changes in the brain in Alzheimer’s disease. Nat Commun 14, 4466, (2023).

24 Yarbro, J. M. et al. Human and mouse proteomics reveals the shared pathways in Alzheimer’s disease and delayed protein turnover in the amyloidome. Nat Commun 16, 1533, (2025).

25 Li, W., et al. An Extensive Atlas of Proteome and Phosphoproteome Turnover Across Mouse Tissues and Brain Regions. bioRxiv, (2024).

26 Liao, L. et al. Proteomic characterization of postmortem amyloid plaques isolated by laser capture microdissection. J. Biol. Chem. 279, 37061–37068, (2004).

27 Xiong, F., Ge, W. & Ma, C. Quantitative proteomics reveals distinct composition of amyloid plaques in Alzheimer’s disease. Alzheimers Dement 15, 429–440, (2019).

28 Drummond, E. et al. The amyloid plaque proteome in early onset Alzheimer’s disease and Down syndrome. Acta Neuropathol Commun 10, 53, (2022).

29 Rahman, M. M. & Lendel, C. Extracellular protein components of amyloid plaques and their roles in Alzheimer’s disease pathology. Mol Neurodegener 16, 59, (2021).

30 Ross-Munro, E. et al. Midkine: The Who, What, Where, and When of a Promising Neurotrophic Therapy for Perinatal Brain Injury. Front Neurol 11, 568814, (2020).

31 Yasuhara, O. et al. Midkine, a novel neurotrophic factor, is present in senile plaques of Alzheimer disease. Biochem Biophys Res Commun 192, 246–251, (1993).

32 Muramatsu, H. et al. Midkine as a factor to counteract the deposition of amyloid beta-peptide plaques: in vitro analysis and examination in knockout mice. Int Arch Med 4, 1, (2011).

33 Oakley, H. et al. Intraneuronal beta-amyloid aggregates, neurodegeneration, and neuron loss in transgenic mice with five familial Alzheimer’s disease mutations: potential factors in amyloid plaque formation. J Neurosci 26, 10129–10140, (2006).

34 LeVine, H., 3rd. Thioflavine T interaction with synthetic Alzheimer’s disease beta-amyloid peptides: detection of amyloid aggregation in solution. Protein Sci 2, 404–410, (1993).

35 Meisl, G. et al. Molecular mechanisms of protein aggregation from global fitting of kinetic models. Nat Protoc 11, 252–272, (2016).

36 Yang, Y. et al. Cryo-EM structures of amyloid-beta 42 filaments from human brains. Science 375, 167–172, (2022).

37 Kollmer, M. et al. Cryo-EM structure and polymorphism of Abeta amyloid fibrils purified from Alzheimer’s brain tissue. Nat Commun 10, 4760, (2019).

38 Lu, J. X. et al. Molecular structure of beta-amyloid fibrils in Alzheimer’s disease brain tissue. Cell 154, 1257–1268, (2013).

39 Sawaya, M. R., Hughes, M. P., Rodriguez, J. A., Riek, R. & Eisenberg, D. S. The expanding amyloid family: Structure, stability, function, and pathogenesis. Cell 184, 4857–4873, (2021).

40 Yang, Y. et al. Cryo-EM structures of amyloid-beta filaments with the Arctic mutation (E22G) from human and mouse brains. Acta Neuropathol 145, 325–333, (2023).

41 Evans, R., O’Neill, M., Pritzel, A., Antropova, N., Senior, A., Green, T., … & Hassabis, D.. Protein complex prediction with AlphaFold-Multimer. biorxiv, DOI: 10.1101/2021.1110.1104.463034, (2022).

42 Roche, J., Shen, Y., Lee, J. H., Ying, J. & Bax, A. Monomeric Aβ(1-40) and Aβ(1-42) Peptides in Solution Adopt Very Similar Ramachandran Map Distributions That Closely Resemble Random Coil. Biochemistry 55, 762–775, (2016).

43 Wallin, C. et al. The Neuronal Tau Protein Blocks in Vitro Fibrillation of the Amyloid-β (Aβ) Peptide at the Oligomeric Stage. J Am Chem Soc 140, 8138–8146, (2018).

44 Yao, Z. et al. A transcriptomic and epigenomic cell atlas of the mouse primary motor cortex. Nature 598, 103–110, (2021).

45 Styren, S. D., Hamilton, R. L., Styren, G. C. & Klunk, W. E. X-34, a fluorescent derivative of Congo red: a novel histochemical stain for Alzheimer’s disease pathology. J Histochem Cytochem 48, 1223–1232, (2000).

46 Ito, D. et al. Microglia-specific localisation of a novel calcium binding protein, Iba1. Brain Res Mol Brain Res 57, 1–9, (1998).

47 Schwabenland, M. et al. Analyzing microglial phenotypes across neuropathologies: a practical guide. Acta Neuropathol 142, 923–936, (2021).

48 Bai, B. et al. Deep Profiling of Proteome and Phosphoproteome by Isobaric Labeling, Extensive Liquid Chromatography, and Mass Spectrometry. Methods Enzymol 585, 377–395, (2017).

49 Wang, Z. et al. 27-plex Tandem Mass Tag Mass Spectrometry for Profiling Brain Proteome in Alzheimer’s Disease. Anal Chem 92, 7162–7170, (2020).

50 Niu, M. et al. Extensive Peptide Fractionation and y1 Ion-Based Interference Detection Method for Enabling Accurate Quantification by Isobaric Labeling and Mass Spectrometry. Anal Chem 89, 2956–2963, (2017).

51 Langfelder, P. & Horvath, S. WGCNA: an R package for weighted correlation network analysis. BMC Bioinformatics 9, 559, (2008).

52 Sousa, C. et al. Single-cell transcriptomics reveals distinct inflammation-induced microglia signatures. EMBO Rep 19, (2018).

53 Zhang, Y. et al. Purification and Characterization of Progenitor and Mature Human Astrocytes Reveals Transcriptional and Functional Differences with Mouse. Neuron 89, 37–53, (2016).

54 Saha, O. et al. The Alzheimer’s disease risk gene BIN1 regulates activity-dependent gene expression in human-induced glutamatergic neurons. Molecular psychiatry, (2024).

55 Voskobiynyk, Y. et al. Alzheimer’s disease risk gene BIN1 induces Tau-dependent network hyperexcitability. Elife 9, (2020).

56 Arvidsson, U., Riedl, M., Elde, R. & Meister, B. Vesicular acetylcholine transporter (VAChT) protein: a novel and unique marker for cholinergic neurons in the central and peripheral nervous systems. J Comp Neurol 378, 454–467, (1997).

57 Rasmussen, N. L., Kournoutis, A., Lamark, T. & Johansen, T. NBR1: The archetypal selective autophagy receptor. J Cell Biol 221, (2022).

58 Cohen, S. I. et al. Proliferation of amyloid-β42 aggregates occurs through a secondary nucleation mechanism. Proc Natl Acad Sci U S A 110, 9758–9763, (2013).

59 Lee, J., Culyba, E. K., Powers, E. T. & Kelly, J. W. Amyloid-β forms fibrils by nucleated conformational conversion of oligomers. Nature chemical biology 7, 602–609, (2011).

60 Belsare, K. D. et al. Soluble TREM2 inhibits secondary nucleation of Aβ fibrillization and enhances cellular uptake of fibrillar Aβ. Proc Natl Acad Sci U S A 119, (2022).

61 Gao, Y., Schedin-Weiss, S. & Tjernberg, L. O. A closer look at pathogenic amyloid-β in Alzheimer’s disease using cryo-electron microscopy: a narrative review. Advanced Technology in Neuroscience 1, 177–187, (2024).

62 Maeda, N. et al. A receptor-like protein-tyrosine phosphatase PTPzeta/RPTPbeta binds a heparin-binding growth factor midkine. Involvement of arginine 78 of midkine in the high affinity binding to PTPzeta. J Biol Chem 274, 12474–12479, (1999).

63 Muramatsu, H. et al. LDL receptor-related protein as a component of the midkine receptor. Biochem Biophys Res Commun 270, 936–941, (2000).

64 Muramatsu, H. et al. alpha4beta1- and alpha6beta1-integrins are functional receptors for midkine, a heparin-binding growth factor. J Cell Sci 117, 5405–5415, (2004).

65 Ichihara-Tanaka, K., Oohira, A., Rumsby, M. & Muramatsu, T. Neuroglycan C is a novel midkine receptor involved in process elongation of oligodendroglial precursor-like cells. J Biol Chem 281, 30857–30864, (2006).

66 Stoica, G. E. et al. Midkine binds to anaplastic lymphoma kinase (ALK) and acts as a growth factor for different cell types. J Biol Chem 277, 35990–35998, (2002).

67 Huang, Y. et al. Midkine induces epithelial-mesenchymal transition through Notch2/Jak2-Stat3 signaling in human keratinocytes. Cell Cycle 7, 1613–1622, (2008).

68 Yamazaki, Y., Zhao, N., Caulfield, T. R., Liu, C. C. & Bu, G. Apolipoprotein E and Alzheimer disease: pathobiology and targeting strategies. Nat Rev Neurol 15, 501–518, (2019).

69 Levites, Y. et al. Integrative proteomics identifies a conserved Abeta amyloid responsome, novel plaque proteins, and pathology modifiers in Alzheimer’s disease. Cell Rep Med 5, 101669, (2024).

70 Chishti, M. A. et al. Early-onset amyloid deposition and cognitive deficits in transgenic mice expressing a double mutant form of amyloid precursor protein 695. J Biol Chem 276, 21562–21570, (2001).

71 Beach, T. G. et al. Arizona Study of Aging and Neurodegenerative Disorders and Brain and Body Donation Program. Neuropathology 35, 354–389, (2015).

72 Connelly, J. P. & Pruett-Miller, S. M. CRIS.py: A Versatile and High-throughput Analysis Program for CRISPR-based Genome Editing. Sci Rep 9, 4194, (2019).

73 Narina, S., Connelly, J. P. & Pruett-Miller, S. M. High-Throughput Analysis of CRISPR- Cas9 Editing Outcomes in Cell and Animal Models Using CRIS.py. Methods Mol Biol 2631, 155–182, (2023).

74 Chen, Z. L. et al. A high-speed search engine pLink 2 with systematic evaluation for proteome-scale identification of cross-linked peptides. Nat Commun 10, 3404, (2019).

75 Yang, Y. et al. Cryo-EM structures of amyloid-β 42 filaments from human brains. Science 375, 167–172, (2022).

76 Delaglio, F. et al. NMRPipe: a multidimensional spectral processing system based on UNIX pipes. J Biomol NMR 6, 277–293, (1995).

77 Lee, W., Tonelli, M. & Markley, J. L. NMRFAM-SPARKY: enhanced software for biomolecular NMR spectroscopy. Bioinformatics 31, 1325–1327, (2015).

78 Schanda, P., Kupce, E. & Brutscher, B. SOFAST-HMQC experiments for recording two-dimensional heteronuclear correlation spectra of proteins within a few seconds. J Biomol NMR 33, 199–211, (2005).

79 Bankhead, P. et al. QuPath: Open source software for digital pathology image analysis. Sci Rep 7, 16878, (2017).

80 Xu, P., Duong, D. M. & Peng, J. Systematical Optimization of Reverse-Phase Chromatography for Shotgun Proteomics. J Proteome Res 8, 3944–3950, (2009).

81 Wang, H. et al. Systematic optimization of long gradient chromatography mass spectrometry for deep analysis of brain proteome. J Proteome Res 14, 829–838, (2015).

82 Wang, X. et al. JUMP: a tag-based database search tool for peptide identification with high sensitivity and accuracy. Mol Cell Proteomics 13, 3663–3673, (2014).

83 Peng, J., Elias, J. E., Thoreen, C. C., Licklider, L. J. & Gygi, S. P. Evaluation of multidimensional chromatography coupled with tandem mass spectrometry (LC/LC- MS/MS) for large-scale protein analysis: the yeast proteome. J Proteome Res 2, 43–50, (2003).

84 Elias, J. E. & Gygi, S. P. Target-decoy search strategy for increased confidence in large-scale protein identifications by mass spectrometry. Nat Methods 4, 207–214, (2007).

85 Zhang, B. & Horvath, S. A general framework for weighted gene co-expression network analysis. Stat Appl Genet Mol Biol 4, Article17, (2005).

86 Yu, G., Wang, L. G., Han, Y. & He, Q. Y. clusterProfiler: an R package for comparing biological themes among gene clusters. OMICS 16, 284–287, (2012).

87 Ritchie, M. E. et al. limma powers differential expression analyses for RNA-sequencing and microarray studies. Nucleic Acids Res 43, e47, (2015).

88 Mertens, B. J. A. Transformation, Normalization, and Batch Effect in the Analysis of Mass Spectrometry Data for Omics Studies. Statistical Analysis of Proteomics, Metabolomics, and Lipidomics Data Using Mass Spectrometry, 1–21, (2016).

